# Repetitive Neuronal Activation Regulates Cellular Maturation State via Nuclear Reprogramming

**DOI:** 10.1101/2025.05.02.651848

**Authors:** Tomoyuki Murano, Hideo Hagihara, Katsunori Tajinda, Keizo Takao, Yoshihiro Takamiya, Kaoru Katoh, Alfred J. Robison, Mitsuyuki Matsumoto, Masakazu Namihira, Tsuyoshi Miyakawa

## Abstract

Neural stimulation, such as electroconvulsive therapy (ECT) and repetitive transcranial magnetic stimulation (rTMS), are highly effective clinical interventions for a broad spectrum of psychiatric disorders, including depression and schizophrenia. However, their mechanism of action at the cellular level remains poorly understood. Here, we modelled ECT with repeated optogenetic neuronal stimulation in the mouse dentate gyrus, and observed ECT-like behavioral effects, including decreased depression-like behavior and increased locomotor activity. At the cellular level, we found dematuration to a long-term stable state, persisting for more than one month, defined by changes in nuclear structure, gene expression patterns resembling the G_2_/M phase of the cell cycle, and altered neural coding of navigational information. Moreover, knockout of the G_2_/M master regulator Cyclin B rescued some of behavioral and cellular effects. These findings demonstrate that ECT-like brain stimulation triggers plasticity of the cellular state, revealing a form of stimulus-regulated nuclear reprogramming with potential clinical utility.

## Introduction

In the adult mammalian brain, strong synchronous neural activity typically has toxic effects on neuronal function and viability. Cumulative electrical activity of sufficiently high amplitude, frequency, and/or duration interferes with cellular metabolism, flooding cells with excess calcium and resulting in metabolic failure and death^1,2^. High-intensity neural activity also results in maladaptive neuronal changes, such as dendritic spine loss^3,4^, abnormal neurite arborization^4,5^, and toxic excitability^6^. Previous studies have identified an intermediate state of senescence after overactivation, characterized by the dematuration of adult neurons to an immature-like state, including global de-differentiation of gene expression patterns, membrane excitability, and neural plasticity^7–11^. Similar neuronal phenotypes have been observed in genetically engineered mice that exhibit abnormal behaviors mirroring common neuropsychiatric disorders, such as schizophrenia, intellectual disability, and bipolar disorder^7,12–16^. Our recent study investigated the levels of lactate, a surrogate marker of neural excitation, in the brains mouse models relevant to neuropsychiatric disorders, and found all mouse models with immature brain phenotypes investigated in this study showed significant increases in lactate levels, suggesting a potential link between cellular hyperexcitability and the immature phenotype^17^. Additionally, immature-like gene expression patterns have been identified in the brain tissues of human patients with neuropsychiatric disorders, such as epilepsy, Alzheimer’s disease, and schizophrenia^11,18–23^, where neural hyperexcitability is a known or suspected etiological contribution.

In marked contrast to the detrimental effects of high-intensity neural activity, there are reported beneficial effects of moderate electrical activity on neural circuit function. Neural activity is essential for brain development, learning and memory^24,25^, including enhancements in synaptic weight^26,27^, neuronal excitability^28,29^, and gene expression^30,31^. An interesting clinical application of moderate activity is the use of electrical brain stimulation technologies such as electroconvulsive therapy (ECT), as a class of methods for ameliorating negative symptoms in patients with major depression^32,33^. ECT promotes neuroplastic changes, such as increased adult neurogenesis in the dentate gyrus and hippocampal volume enlargement^34–36^ with 6–12 repeated stimulations commonly used for long-term positive clinical outcomes^32,33^. Electroconvulsive stimulation (ECS), the animal equivalent of ECT, also induces molecular and electrophysiological changes, such as enhanced neurogenesis and altered synaptic transmission^9,14,37,38^. Similar changes occur with chronic administration of fluoxetine, a selective serotonin reuptake inhibitor (SSRI)^9,39,40^. Interestingly, repetitive ECS and chronic fluoxetine administration can induce cellular dematuration, suggesting that unspecified cellular plasticity may underpin their antidepressant effects^9,39–41^ Likewise, other noninvasive brain stimulation methods, such as repetitive transcranial magnetic stimulation (rTMS) and transcranial direct current stimulation (tDCS), can induce neuroplastic changes in the brain and have promising therapeutic potential ^42,43^. However, despite their common use, the specific cellular mechanisms engaged by these brain stimulation therapies, including clinical ECT, rTMS, and tDCS, remain poorly understood.

In this study, we aimed to elucidate the primary cellular mechanisms by which brain stimulation technologies like ECT cause cellular dematuration. We applied an ECT-like optogenetic stimulation to granule cells of the hippocampal dentate gyrus, a key brain area for ECT, to produce clinically relevant behavioral effects. This stimulation led to an immature-like gene expression profile in the brain of mice, and a similar patten was observed in the post-mortem brains of patients with mood disorders. Our analysis also revealed that repetitive stimulations induce G_2_/M cell cycle re-entry and nuclear structural plasticity with epigenomic and morphonuclear signatures. These findings reveal a unique immature cellular state induced by ECT-like neuronal activation, which has long-term stability, and a mechanism of activity-dependent nuclear reprogramming, offering mechanistic insight into ECT effects and related applications.

## Results

### 1. Neuronal activations trigger an immature cellular state in DG granule cells

We challenged neurons *in vivo* with a stimulation protocol designed to mimic human ECT, in which seizure duration is a critical determinant of therapeutic efficacy and is therefore tightly controlled in clinical settings. Conventional methods, such as seizure-inducing drugs and animal ECS, do not allow precise control over seizure duration, leading to variability in cellular and network-level effects. To overcome this limitation, we used mice expressing channelrhodopsin-2 (ChR2) in granule cells (GCs) of the dentate gyrus (DG) (POMC-Cre::ChR2-EYFP mice) and applied optogenetic stimulation, termed Repetitive Optogenetic Stimulation (REPOPS), enabling precise control over the frequency and duration of stimulation (Fig.1a and Extended Data Fig. 1a). This optogenetic approach allowed us to establish a precise experimental system to study cellular effects confined to a specific target cell population. We employed a moderate stimulation protocol that did not cause cell death (Extended Data Fig. 1b–d).

REPOPS was performed either three or ten times consecutively (Fig. 1a). The expression of calbindin (CB), a maturation marker of GCs, was reduced 24 hours after REPOPS for three days, but returned to a level that was not significantly different from the pre-stimulus levels within two weeks (Fig. 1b). In contrast, REPOPS for 10 days significantly decreased CB expression at 24 hours and two weeks after the last stimulation (Fig. 1b). The decrease in CB was inhibited by the administration of Ca^2+^ blockers (Extended Data Fig. 2a– c), suggesting that Ca^2+^ influx is crucial for REPOPS-induced dematuration. RNA-seq analysis showed that the number of genes with significant changes in expression (*P* < 0.05, fold change > 1.2; vs. No Stim group) decreased over time after 3-day REPOPS, but remained unchanged after 10-day REPOPS (Fig. 1c). Evidently, neuronal activation repeated a few times induces short-term transcriptome changes that spontaneously revert to the original state; however, when neuronal activation is chronically repeated, the changes are sustained for longer periods. To examine whether REPOPS induces an immature-like gene expression pattern at the transcriptome level, we assessed the similarity between gene expression patterns of optogenetically-stimulated DG and those of normal infant mice (GSE113727^11^) by the Running Fisher test^44^, a rank-based nonparametric algorithm that evaluates the degree of overlap between two gene groups implemented in BaseSpace (Illumina, Cupertino, CA, USA). There was a striking degree of similarity between the datasets (Overlap *P* = 3.9 × 10^−52^) with the same directional changes in expression (Fig. 1d). Collectively, these results indicate that REPOPS for 10 consecutive days induces long-lasting immature-like gene expression patterns in the DG, whereas REPOPS for three days does not.

**Figure 1.**
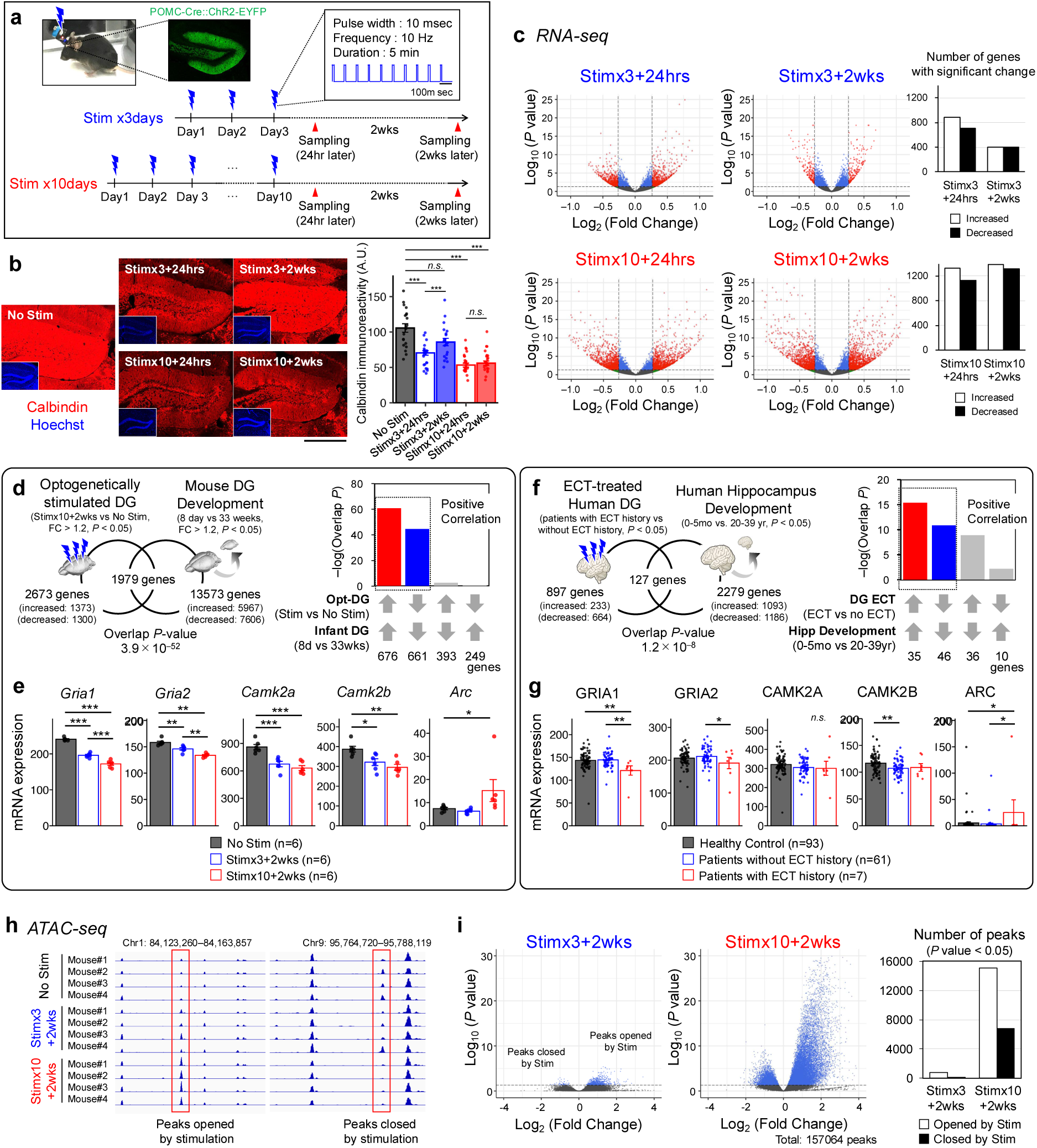
REPOPS induces long-term changes in gene expression, chromatin accessibility, and locomotor activity in mice. **a,** Experimental design. POMC-Cre::ChR2-EYFP mice express ChR2 in GCs of the DG. Blue light was delivered via a wireless LED device. Optogenetic stimulation (10-msec pulse at 10 Hz for 5 min/day) was applied for 3 or 10 consecutive days. Brains were collected 24 hours or two weeks after the last stimulation. **b,** Immunohistostaining of calbindin, a maturation marker of GCs, in dorsal DG. Scale bar, 500 μm. Bar graph shows calbindin immunoreactivity (10–15 sections from 4 mice in each group). Error bars represent the Mean ± s.e.m. One-way ANOVA followed by Bonferroni correction for multiple comparisons, *F*_(4,106)_ = 24.9, *P* = 1.5 × 10^−14^, \*\*\**P* < 0.001. **c,** RNA-Seq. Volcano plots show fold change and *P* values distribution after differential transcript abundance analysis using DESeq2. All stimulation groups were compared with the No Stim group (n = 6 mice per group). Genes with significantly differential expression (*P* < 0.05; fold change > 1.2) are highlighted in red. Bar graphs indicate number of genes with significant changes (i.e. red dots in Volcano plots). **d,** Overlap between genes altered by REPOPS (Stim×10+2wks vs. No Stim) and during normal development of the mouse DG (P8 infants vs. 33-week-old adults; GSE113727). Bar graphs show–log_10_ of overlap *P-*values for genes upregulated or downregulated by each condition. The Bonferroni correction was used to adjust the significance level according to the number of dataset pairs. **e,** mRNA expression levels (FPKM) of representative synapse-related genes. Mean ± s.e.m. (n = 6 mice/group). One-way ANOVA followed by Bonferroni correction for multiple comparisons; \**P* < 0.05, \*\**P* < 0.01, \*\*\**P* < 0.001. **f,** Overlap between genes altered in DG of human patients with bipolar disorder and major depressive disorder (with a history of ECT vs. without ECT; SRP241159^45^) and those altered during human hippocampal development (0-5 month infants vs. 20-39-year-old adults; GSE25219^46^). **g,** mRNA expression levels (FPKM) of representative synapse-related genes in post-mortem DG from healthy control subject (n=93), patients without ECT history (n=61), and patients with ECT history (n=7). **i** ATAC-seq. Integrative Genomics Viewer (IGV) visualization of representative gained-open and gained-closed chromatin profiling coverage. **j,** Volcano plot showing the fold change and *P* value distribution after differential analysis of chromatin accessibility using DESeq2. Stim×3+2wks and Stim×10+2wks groups were compared with the No Stim group (n = 4 mice/group). Bar graph shows number of peaks with a significant change (*P* < 0.05).

As REPOPS was designed to model the effects of human ECT, we next asked whether ECT in human patients also induces an immature-like gene expression pattern. We reanalyzed RNA-seq data from the post-mortem DG of bipolar disorder and major depressive disorder patients with and without ECT (SRP241159^45^). Genes differentially expressed between ECT-treated and non-ECT-treated patients were compared with those changing during typical hippocampal development (0–5 months vs. 20–39 years; GSE25219^46^). This analysis revealed a significant overlap between the two datasets (Overlap *P* = 1.2 × 10⁻^8^), with consistent directional changes (Fig. 1f), suggesting that ECT induces an immature-like gene expression pattern in the DG of human patients, in line with the pattern observed in REPOPS. Notably, both REPOPS and human ECT induced long-term alterations in the expression of synapse-related genes: REPOPS significantly downregulated *Gria1*, *Gria2*, *Camk2a*, and *Camk2b*, and upregulated *Arc* (Fig. 1e), while ECT-treated patients exhibited similar alterations in GRIA1, GRIA2, and ARC compared with non-ECT-treated patients (Fig. 1g), suggesting a shift toward an immature synaptic state.

### 2. REPOPS regulates the long-term plasticity of genomic structure

Changes in gene expression are often accompanied by alterations in the chromatin structure of the associated genomic regions^47^. To investigate whether and how neural overactivation alters the chromatin accessibility of neurons, we performed ATAC-seq on optogenetically stimulated DG tissue (Fig. 1h). Volcano plots showing changes in chromatin accessibility are shown in Fig. 1i. We found significant changes in chromatin accessibility in 806 chromatin regions (748 opened and 58 closed) and 21,853 chromatin regions (15,098 opened and 6,755 closed) in the Stim×3+2wks and Stim×10+2wks groups, respectively (Fig. 1i). These results indicate that repeated neuronal activation for 10 consecutive days, but not for 3 days, induced widespread and persistent reorganization of three-dimensional genome structure, along with concordant global transcriptome changes.

### 3. REPOPS initiates long-term anti-depressive behavioral changes in mice

To assess whether ECT-like neuronal activation could alter anti-depressive behavior, we first performed an open field (OF) test (Fig. 2a, 2b). Locomotor activity was recorded for 30 min each day, and the dorsal DG was optogenetically stimulated for 5 min (20–25 min after the start) on each stimulation day. As previously reported^48^, mice showed an increase in locomotor activity immediately after stimulation (a1–a3 in Fig. 2a). Interestingly, on the day after the first stimulation, mice in both Stim×3 and Stim×10 groups exhibited a significant increase in distance traveled during the first 15 min compared to the No Stim group (*P* = 0.0134, Day 2 in Stim×10 group; *P* = 8.05 × 10^−4^, Day 9 in Stim×3 group; Fig. 2b). In the Stim×3 group, this increase peaked on Day 9 and returned to control levels within two weeks (blue line in Fig. 2b), whereas in the Stim×10 group, it declined gradually from the last stimulation day but remained significantly elevated two weeks after the last stimulation (Day 24, *P* = 0.00362; red line in Fig. 2b). Thus, neuronal activation of the DG repeated for 10 days induced hyperlocomotor activity persisting for at least two weeks, whereas an increase in locomotor activity after neuronal activation for three days was transient. We also conducted 24-hour activity monitoring in the home cage (Fig. 2c). The distance traveled by mice in the Stim×10 group gradually increased after the initial stimulation and was significantly higher than that in the No Stim and Stim×3 groups (*P* < 0.05, Days 7–36; Fig. 2c). Mice in the Stim×3 group did not show any significant increase in locomotor activity in the home cage. These results indicate that repeated neuronal activation causes a significant increase in baseline locomotor activity, which persists for more than one month. Depression-related behavior was assessed with the tail suspension test (Fig. 2d). Mice in the Stim×10 group showed significantly less immobility compared to those in the No Stim (*P* = 0.0094) and Stim×3 groups (*P* = 0.0198). A similar trend was observed in the forced swim test (Fig. 2e), where Stim×10 mice showed reduced immobility compared to the No Stim group (*P* = 0.0304) and Stim×3 group (*P* = 0.0770). These results suggest that repeated neuronal activation mimicking ECT reduces depression-like behavior in mice (for other behavioral tests, see also Extended Data Fig. 3).

**Figure 2.**
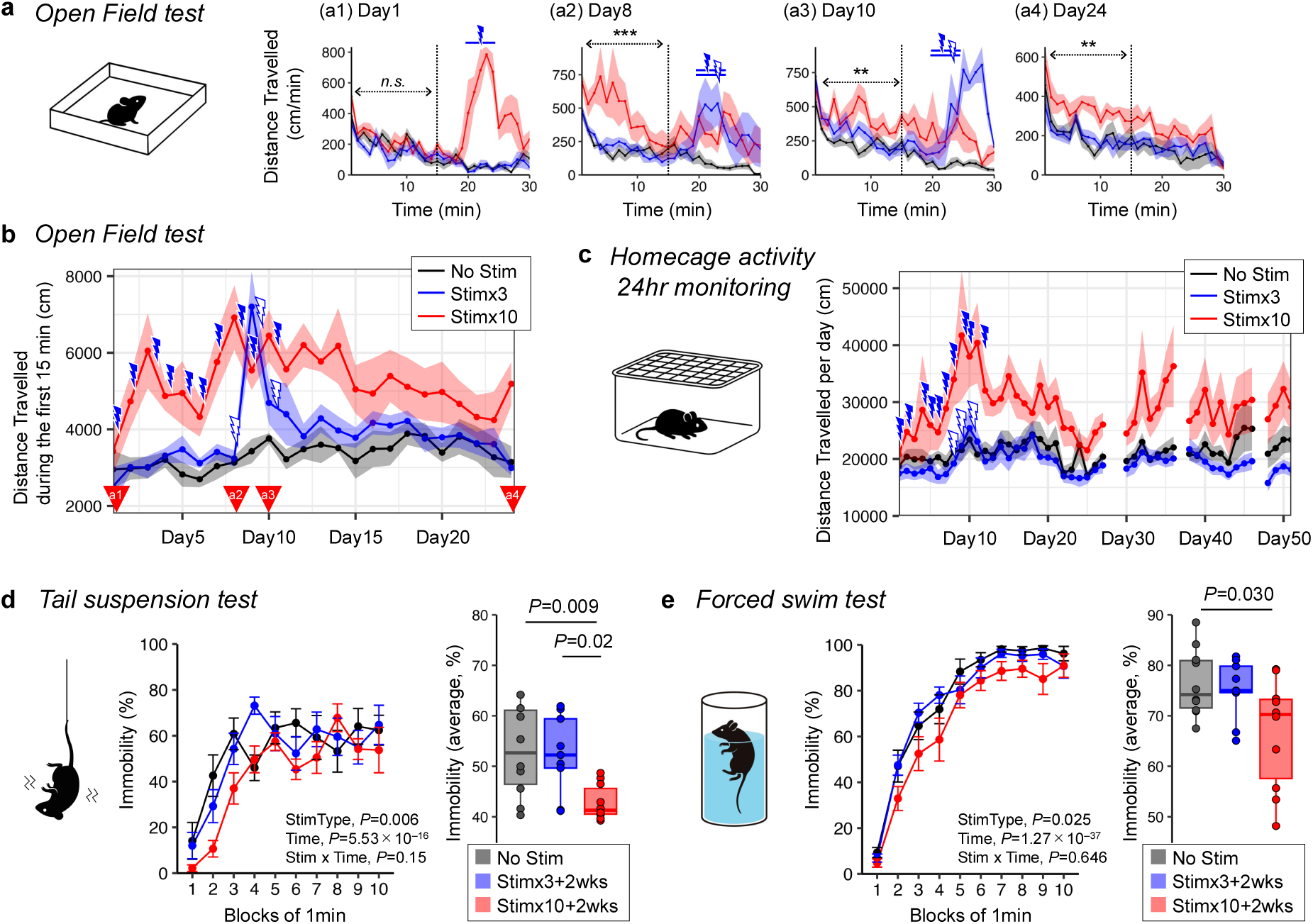
REPOPS induces anti-depressive behaviors. **a,** Distance traveled in the open field during the first 30 min on Day 1 (a1), Day 8 (a2), Day 10 (a3), and Day 24 (a4), corresponding to time points in **b** (red arrowheads). Solid lines show group means (n = 6 mice/group); semitransparent area indicates ± s.e.m. (black, No Stim; blue, Stim×3+2wks; red, Stim×10+2wks). Optogenetic stimulation was introduced for 5 min (20– 25 min after the start) on Day 1, Day 8, and Day 10. Asterisk indicates Stim×10+2wks vs. No Stim; One-way ANOVA with Bonferroni correction; \**P* < 0.05, \*\**P* < 0.01, \*\*\**P* < 0.001. **b,** Total distance traveled in the first 15 min of the open field test from Day 1 to Day 24. Two-way repeated measures ANOVA; Stim type, *F*_(2, 14)_ = 9.96, *P* = 2.0×10^−3^; Day, *F*_(23, 322)_ = 5.08, *P* = 4.63×10^−12^. Stim type × Day, *F*_(46, 322)_ = 2.58, *P* = 7.81×10^−7^. **c,** 24-hour locomotor activity in the home cage. Daily average distance traveled is shown (n = 10, 8, and 10 mice for No Stim, Stim×3, and Stim×10 groups, respectively). Data missing on Day 28, 29, 37 and 47 due to computer system error. *P* < 0.05 on Day 7–8, 10–16, 20, 22–23, 25–30, 33–36 (Stim×10 vs. No Stim). **d,** Immobility (%) in tail suspension test. Values are the Mean ± s.e.m. Two-way repeated-measures ANOVA results shown in panel. n = 10, 9, and 10 mice. Box plot summarizes average immobility over 10 min. One-way ANOVA with Bonferroni correction; \**P* < 0.05, \*\**P* < 0.01, \*\*\**P* < 0.001. **e,** Same as **d**, for immobility (%) in the forced swim test.

### 4. Alterations of neural coding of navigational information

To investigate whether and how neural overactivation affects circuit function, we performed *in vivo* Ca^2+^ imaging under freely moving conditions combined with optogenetic stimulation (Fig. 3a, 3b). On each stimulation day, we recorded DG neuron activity and locomotor activity in the OF during the first 30 min, followed by 5 min of optogenetic stimulation (590–650 nm, 20 msec pulse, 10 Hz; Fig. 3c). The distance traveled during the first 30 min in the Stim×10 group gradually increased and was significantly greater than in the No Stim group two weeks after the last stimulation (*P* = 0.0358; Fig. 3d). However, the mean Ca^2+^ transient rate did not differ significantly between these groups throughout the experiment (Fig. 3e).

**Figure 3.**
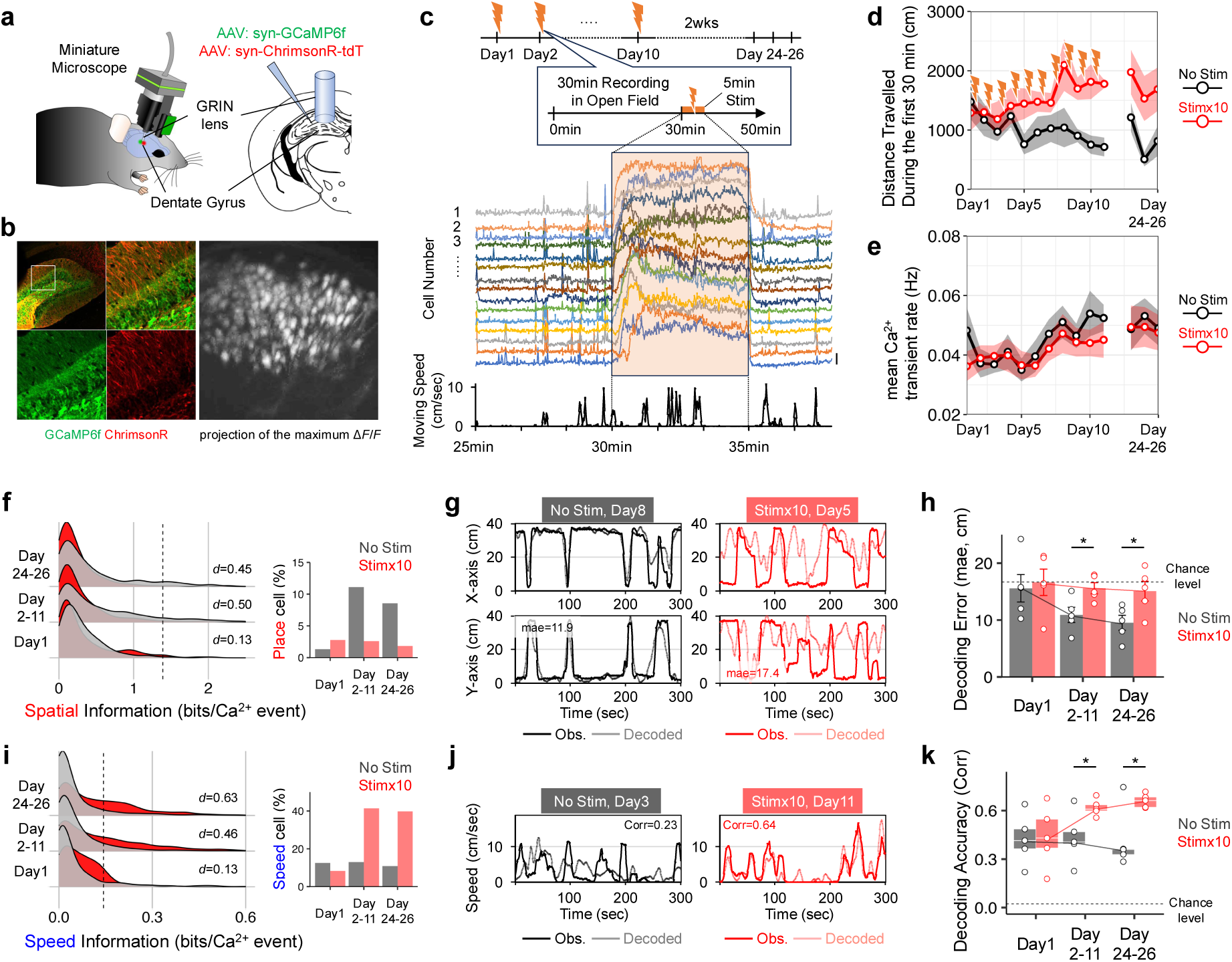
REPOPS mediates long-term changes in neural coding of navigational information. **a,** Simultaneous *in vivo* calcium imaging and optogenetic stimulation under free-moving conditions. AAV-syn-GCaMP6f and AAV-syn-ChrimsonR were co-injected into the DG, followed by GRIN-lens implantation. An integrated miniature microscope (nVoke; Inscopix) enabled large-scale cellular-resolution Ca^2+^ imaging and optogenetic stimulation. **b,** Left: Expression of GCaMP6f and ChrimsonR in the DG. Right: Max projection of relative fluorescence change (Δ*F*/*F*) of Ca^2+^ transients from all imaging frames during the first 30 min, showing representative active neurons. **c,** Top: Experimental design of Ca^2+^ imaging combined with optogenetic stimulation during the open field test. Each 50-min session consisted of 30-min Ca^2+^ imaging (used for the following analysis), 5-min optogenetic stimulation, and 15-min post-stimulation recording. This was repeated for 10 days. No light was introduced to mice in the No Stim group. Bottom: Δ*F*/*F* traces from 15 representative neurons (scale bar, 10% Δ*F*/*F*). Data from 25 to 38 min of a 50-min session are shown. **d,** Distance traveled during the first 30 min **e,** Average Ca^2+^ transient rate during the first 30 min. **f,** Distribution of spatial information for the No Stim (grey curve) and Stim×10 groups (red curve). The vertical axis represents the frequency of distribution, with the dotted line indicating the criterion for place cells (top 95% percentile of the shuffled distribution; see Methods section). Bar graph indicates proportion of place cells. **g,** Representative results of position decoding using Ca^2+^ imaging data. The first 30 min was split into two 15-min halves for training and test data for decoding. Black/red lines: observed position; grey/pink lines: decoded position. **h,** Decoding accuracy (mae; mean absolute error, cm). Dotted lines: shuffled control. Two-way repeated measures ANOVA: Stim type, *F*_(1, 8)_ = 6.49, *P* = 0.034; Day, *F*_(2, 16)_ = 2.89, *P* = 0.085; Stim type × Day, *F*_(2, 16)_ = 1.11, *P* = 0.353. Bonferroni correction for multiple comparisons was performed, \**P* < 0.05. **i,** Same as **f**, but for speed information and speed cells. The dotted line indicates the criterion for speed cells (top 99% percentile of the shuffled distribution). **j,** Same as **g**, but for speed decoding. **k,** Same as **h**, but for speed decoding accuracy. Two-way repeated measures ANOVA: Stim type, *F*_(1, 8)_=4.02, *P* = 0.080; Day, *F*_(2, 16)_ = 4.87, *P* = 0.022; Day×Stim type, *F*_(2, 16)_ = 3.98, *P* = 0.039. Bonferroni correction for multiple comparisons was performed, \**P* < 0.05.

Navigational information, such as position and speed, is encoded in the population activity of DG neurons^16,49^. To examine how neural overactivation affects this coding, we computed spatial and speed information for individual DG neurons. In the No Stim group, spatial information increased on Days 2–11 and Days 24–26 compared to Day 1 (Cohen’s *d* = 0.43 and 0.30), suggesting that OF exposure enhanced spatial coding (Fig. 3f). In contrast, the Stim×10 group showed reduced spatial information on Days 2–11 and Days 24–26 compared to the No Stim group (*d* = –0.50 and –0.45; Fig. 3f). Speed information remained unchanged in the No Stim group throughout the experiment (|*d|* < 0.10), but increased in the Stim×10 group on Days 2–11 and Days 24–26, exceeding that in the No Stim group (*d* = 0.46 and *P* = 0.63; Fig. 3i). Thus, repeated neuronal activation reduced spatial coding and enhanced speed-related activity in individual DG neurons for over two weeks.

To examine information encoding at the population level, we predicted mouse position and speed from the population activity of DG neurons using machine learning methods. Representative decoding results are shown in Fig. 3g and 3j. In the No Stim group, position decoding errors decreased on Days 2–11 and Days 24–26 and were significantly smaller than chance (Fig. 3h). In contrast, the Stim×10 group showed no significant change in position decoding accuracy, and decoding errors were significantly greater than in the No Stim group on Days 2–11 and Days 24–26 (*P* = 0.0133 and 0.0335; Fig. 3h). These findings suggest that position information coding was enhanced by OF exposure but impaired by neural overactivation. Speed decoding accuracy did not change significantly during the experiment in the No Stim group, but was increased by REPOPS in the Stim×10 group and was significantly higher than in the No Stim group on Days 2–11 and Days 24–26 (*P* = 0.0363 and 0.0223; Fig. 3k). Thus, repeated neuronal activation reshapes information processing in the DG by suppressing spatial map formation and enhancing speed-related activity, with unchanged overall activity levels. This shift may underlie the behavioral changes observed in optogenetically stimulated mice, such as increased locomotor activity and reduced depression-like behavior.

### 5. REPOPS-driven long-term increases in the expression of cell cycle-related genes

We next explored potential molecular markers associated with REPOPS-induced neuronal alterations by conducting gene ontology analysis of RNA-seq data from optogenetically-stimulated DG. Notably, gene categories significantly altered in the Stim×10+2wks group—but not in the Stim×3+2wks group—included cell cycle–related terms, such as *“cell division”* and *“mitotic cell cycle process”* (Extended Data Fig. 4). Expression levels of representative cell cycle-related genes are shown in Fig. 4a. Genes related to the G_2_/M phase, such as *Ccna2*, *Ccnb1*, *Ccnb2*, and *Cdc25c*, were significantly increased in Stim×10+2wks compared to the No Stim group, whereas genes related to other phases, such as *Ccne1*, *Ccne2*, *Cdk2* (G_1_-S phase), and *Ccnd1*, *Cdk4* (G_1_ phase), showed no significant changes (Fig. 4a). We found that cyclin B, an essential regulator for the transition from G_2_ to M phase, was expressed in most DG neurons in Stim×10+2wks, while it was barely detectable in the No Stim group (Fig. 4b). It was broadly expressed throughout the granule cell layer and was not restricted to the subgranular zone, where newborn neurons are abundant, indicating that the majority of cyclin B expression reflects induction in mature neurons, rather than enhanced neurogenesis.

**Figure 4.**
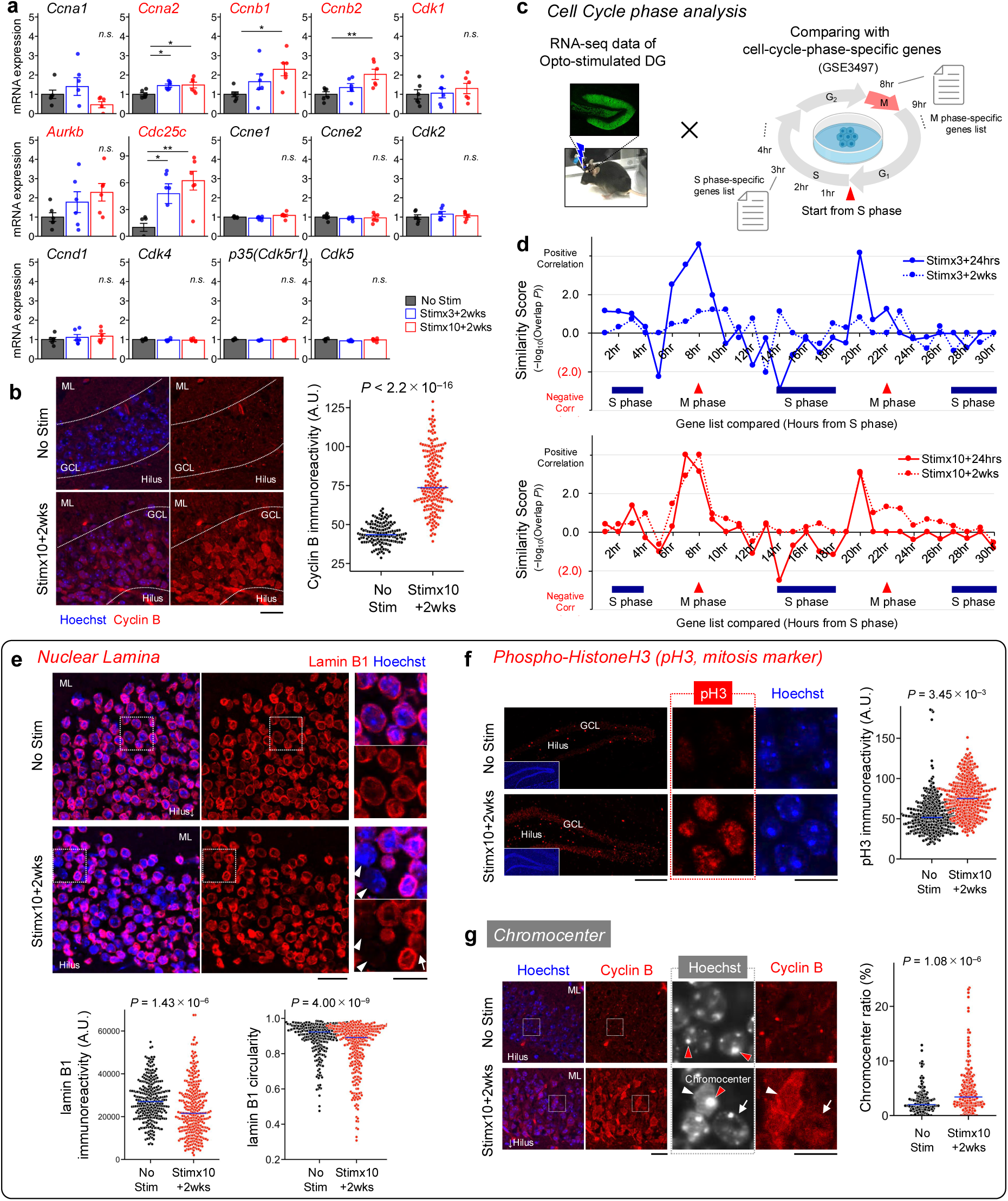
REPOPS confers G_2_/M phase-like gene expression patterns and nuclear structure plasticity. **a,** Expression of representative cell cycle-related genes. mRNA expression levels (FPKM) are normalized to the control group (No Stim). Data are shown as mean ± s.e.m. (n = 6 mice/group). One-way ANOVA with Bonferroni correction; \**P* < 0.05, \*\**P* < 0.01. **b,** Cyclin B immunostaining (red) in the DG. Beeswarm plot shows cyclin B immunoreactivity in individual neurons. Blue bars indicate the median values of 147 neurons (No Stim) and 202 (Stim×10+2wks) neurons from three biologically independent samples. ML, molecular layer; GCL, granule cell layer. **c,** Cell cycle phase analysis. To infer the cell cycle-like status of DG neurons after REPOPS, RNA-seq data from optogenetically stimulated DG (Fig. 1c) were compared with publicly available reference gene expression profiles representing each phase of the cell cycle. These reference profiles were derived from microarray data of HeLa cell sampled at 1-hour intervals (1-30 h) after release from S phase arrest (Whitfield *et al.*, 2002; GSE3497)^50^. **d,** Similarity in gene expression patterns between optogenetically stimulated DG and gene expression profiles representing each phase of the cell cycle. Similarity scores were calculated as –log of the overlap *P* value between RNA-seq data of stimulated DG and gene sets representing each time point of the cell cycle (1–30 hours from the start of S phase). Positive scores indicate concordant expression changes (e.g., both upregulated), while negative scores indicate opposite trends. The positions of the S and M phases on the x-axis reflect cell cycle timing estimated from the HeLa cell data (GSE3497; Whitfield et al., 2002)^50^. **e,** Lamin B1 immunostaining (red) in the DG. Enlarged views of boxed regions are shown at right. Arrows and arrowheads indicate neurons with partial or complete disruption of lamin B1 structure. Scale bars: 20 μm (left), 10 μm (right). Beeswarm plots show immunoreactivity and circularity of lamin B1 in individual neuron (n = 289 and 320 neurons from three independent samples). Blue bars, medians. Welch’s *t*-tests: immunoreactivity, *t*_(589)_ = 4.87, *P* = 1.45 × 10^−6^; circularity, *t*_(547)_ = 5.87, *P* = 7.38 × 10^−9^. **f,** Immunostaining of phospho-histone H3 (pH3) (red) in the DG. Scale bars: 200 μm (left), 10 μm (right). Beeswarm plot shows pH3 signal intensity in individual neurons (n = 415 and 459 neurons from three independent samples). Welch’s *t*-test: *t*_(858)_ = 14.1, *P* < 2.2 × 10^−16^. **g,** DNA counterstaining with Hoechst (blue and white pseudo-color) and cyclin B immunostaining (red) in the DG. Enlarged views of boxed areas are shown on the right. Red arrowheads indicate chromocenters–an aggregation of the heterochromatin domain visible by Hoechst staining. White arrowheads/arrows: neurons with large/small chromocenters and high/low cyclin B expression, respectively. Scale bars: 100 μm (left), 10 μm (right). Beeswarm plot shows the chromocenter ratio (%) per neurons (n = 147 and 202 neurons from three independent samples). Welch’s *t*-test: *t*_(306)_ = 5.46, *P* = 9.38 × 10^−8^.

To assess whether gene expression patterns induced by REPOPS resembled specific cell cycle phases, we used a publicly available microarray dataset of HeLa cells that captures gene expression pattern dynamics throughout the cell cycle (Fig. 4c; GSE3497^50^). In this dataset, genes characteristic of each phase were identified through microarray analysis of HeLa cells sampled at multiple time points across the cell cycle. We compared these cell cycle phase-specific gene lists (1–30 hour from S phase) with RNA-seq data from optogenetically stimulated DG (Fig. 4c) and computed a “similarity score” for each time point, defined as the –log_10_ of the overlap *P* value (Fig. 4d). In Stim×3+24hrs and Stim×10+24hrs, similarity score peaks were observed at 7, 8, and 20 hours from S phase (scores > 3.0; Figs. 4d), corresponding approximately to the G_2_/M phase. In Stim×3+2wks, these peaks declined (scores < 2.0), whereas in the Stim×10+2wks group, they remained elevated (scores > 3.0; Figs. 4d). Although neurons are considered post-mitotic and do not undergo cell cycle progression, these findings suggest that repeated neuronal activation reinstates G_2_/M phase-like transcriptional programs in mature neurons that persist for over two weeks.

### 6. Long-term G_2_/M phase-like epigenomic plasticity in nuclear structure

We investigated whether REPOPS can induce changes in nuclear structure, focusing on three major events that occur during the G_2_/M phase: (i) nuclear lamina disruption, (ii) an increase in the phosphorylation of histone H3 (pH3), and (iii) chromatin condensation.

i. The nuclear lamina is a dense fibrillar network lining the inner nuclear membrane and providing mechanical support for chromatin organization, which disrupts during mitosis due to nuclear division^51,52^. In the No Stim group, most neurons exhibited normal, round-shaped lamin B1, the main component of the nuclear lamina (Fig. 4e and Extended Data Fig. 5a, 5b). In contrast, neurons in the Stim×10+2wks group showed varying degrees of reduced expression and/or disrupted lamin B1 structure, indicating that 10 days of REPOPS induced nuclear lamina disruption (Fig. 4e).
ii. Increased phosphorylated histone H3 (pH3) is another characteristic change during mitosis, which is a known “mitosis marker.” In the No Stim group, most neurons showed minimal immunoreactivity of pH3, whereas the Stim×10+2wks group exhibited a significant increase in pH3 immunoreactivity (Fig. 4f), suggesting that repeated neuronal activation induces mitosis-like epigenetic modifications of chromatin in mature neurons.
iii. The chromocenter is an aggregation of heterochromatin domains, which can be visualized as concentrated dot-like signals in the nucleus by DNA counterstaining (indicated by red arrowheads in Fig. 4g). It is considered a central site for chromatin condensation during mitosis^53,54^. The chromocenter ratio–defined as the ratio of total chromocenter area to the nucleus size–was significantly increased by REPOPS, indicating an enlargement of heterochromatin domains (Fig. 4g). Thus, REPOPS induces a G_2_/M phase-like nuclear states, characterized by nuclear lamina disruption, increased pH3, and chromatin condensation (see also Extended Data Fig. 6a, 6b).

### 7. Cyclin B mediates REPOPS-induced changes in nuclear structure, cellular immaturity, and behavior

To investigate the molecular mechanisms underlying G_2_/M phase-like nuclear alterations, we knocked out the expression of cyclin B–an essential gene for the G_2_/M phase transition–by injecting a mixture of AAV9-SpCas9 and AAV9-sgRNA encoding sequences targeting *Ccnb1* and *Ccnb2* (CyclinB-KO) into the DG (Fig. 5a, 5b). AAV9-sgRNA encoding scrambled sequences (Scram-KO) was used as a control. The REPOPS-induced expression of cyclin B, observed in neurons in the granule cell layer, was effectively eliminated in the CyclinB-KO group (Extended Data Fig. 7b).

**Figure 5.**
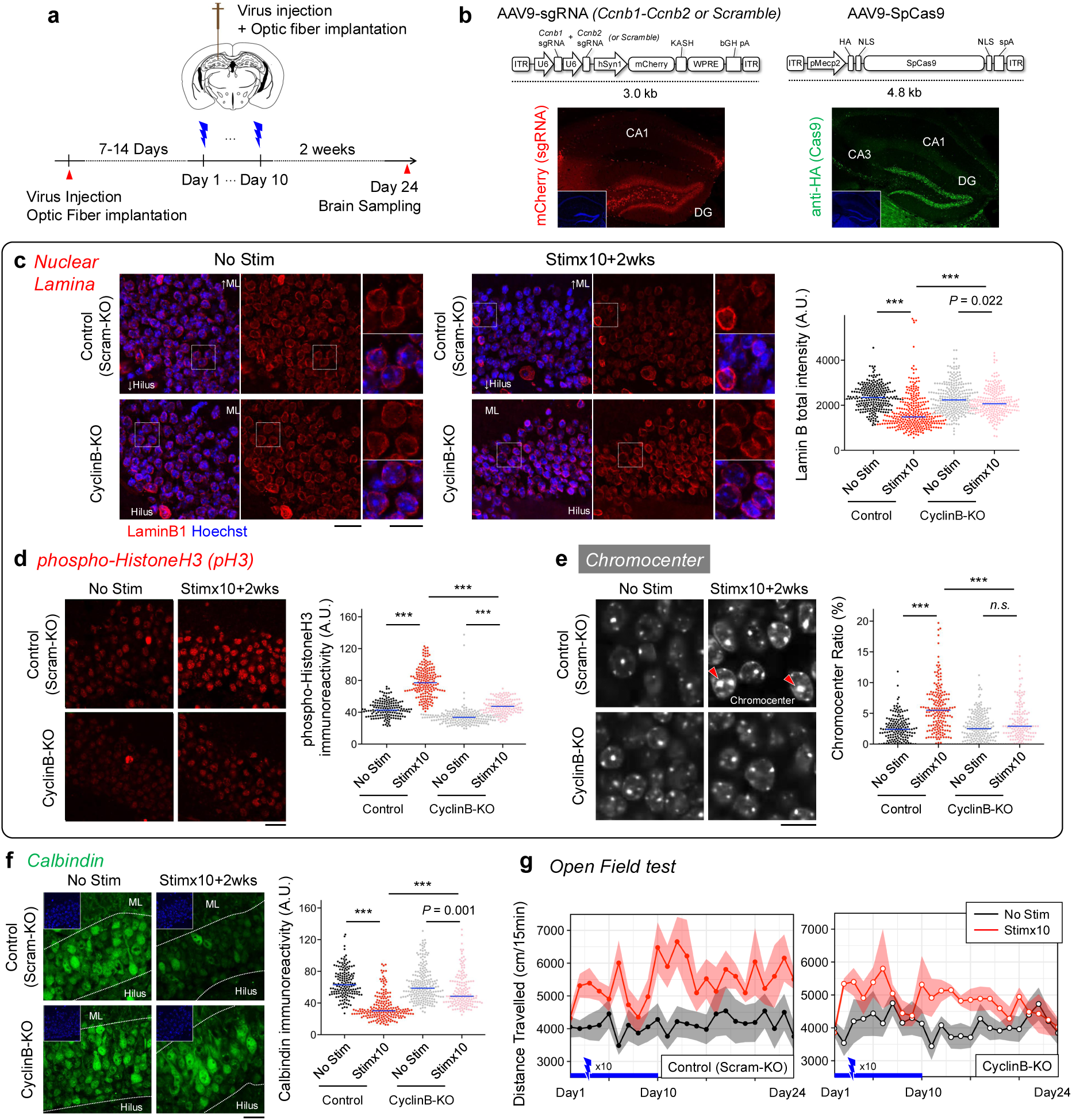
Cyclin B deletion reverses multiple REPOPS-induced phenotypes. **a,** Experimental design. AAV9 vectors encoding Cas9 and sgRNAs targeting *Ccnb1* and *Ccnb2* (CyclinB-KO) or a *Scramble* sequence (Scram-KO) were co-injected into the DG, followed by optic fiber implantation. After 7-14 days, optogenetic stimulation was performed for 10 consecutive days. Brains were sampled 2 weeks after the last stimulation. The open field test was conducted from Day 1 (immediately before the first stimulation) to Day 24 (two weeks after the last stimulation). **b,** Co-expression of sgRNAs and Cas9 in the DG. sgRNAs were co-expressed with mCherry, and Cas9 was tagged with HA. Most neurons in the granule cell layer showed both mCherry and HA immunoreactivity. **c,** Lamin B1 immunostaining in the DG (converted to red for visibility; originally stained with Alexa647). Upper and lower panels show control (Scramble-KO) and *Ccnb1*-*Ccnb2*-KO (CyclinB-KO) mice, respectively. Scale bars: 20 μm (left), 10 μm (right). Beeswarm plot shows lamin B1 immunoreactivity. Red bars, median. One-way ANOVA, *F*_(3, 679)_ = 60.19, *P* < 2.0 × 10^−16^; Bonferroni corrected, \*\*\**P* < 0.001. n = 245, 301, 271, and 246 neurons from three independent samples per group. **d,** pH3 immunostaining in the DG. One-way ANOVA, *F*_(3, 679)_ = 325.7, *P* < 2.0 × 10^−16^; \*\*\**P* < 0.001; 199, 158, 143, and 183 neurons from five independent samples per group. Scale bar, 20 μm. **e,** Chromocenter visualized by Hoechst staining (white, pseudo-color). One-way ANOVA, *F*_(3, 679)_ = 39.1, *P* < 2.0 × 10^−16^; \*\*\**P* < 0.001; 199, 158, 143, and 183 neurons from five independent samples per group. Scale bar, 10 μm. **f,** Calbindin staining in the DG. One-way ANOVA, *F*_(3, 725)_ = 104.3, *P* < 2.0 × 10^−16^; \*\*\**P* < 0.001; 217, 157, 162, and 198 neurons from five biologically independent samples per group. Scale bar, 20 μm. **g,** Open field test: total distance traveled during the first 15 min from Day 1 to Day 24. Left: Scramble-KO; Right: CyclinB-KO. Black and red lines represent No Stim and Stim×10+2wks groups, respectively. Solid lines, group means (n = 7 mice/group); semitransparent areas, ±s.e.m. Two-way repeated measures ANOVA: Groups, *F*_(3, 24)_ = 4.271, *P* = 0.015; Day, *F*_(6, 163)_ = 4.271, *P* = 0.0204; Groups×Day, *F*_(20, 163)_ = 1.09, *P* = 0.36.

Lamin B1 expression was significantly reduced by REPOPS in both Scram-KO and CyclinB-KO groups, but the degree of reduction was significantly smaller in the CyclinB-KO group (Fig. 5c). Similarly, REPOPS significantly increased pH3 signals in both groups, but the degree of increase was significantly reduced in the CyclinB-KO group (Fig. 5d). The chromocenter ratio was significantly increased by REPOPS in the Scram-KO group, but not in the CyclinB-KO group (Fig. 5e). Calbindin expression was significantly decreased in both groups, but the reduction was significantly smaller in the CyclinB-KO group compared to the Scram-KO group (Fig. 5f). In the open field test, REPOPS significantly increased locomotor activity in the Scram-KO group from the day after the first stimulation, which remained significantly greater than that of the No Stim group two weeks after the last stimulation (*P* = 0.0231, Day 24; Fig. 5g). In contrast, in the CyclinB-KO group, locomotor activity showed a transient increase but gradually returned to baseline levels by Day 24 (*P* > 0.99; Fig. 5g). Collectively, these findings demonstrate that cyclin B is a critical molecular mediator linking repeated neuronal activation to G_2_/M phase-like nuclear remodeling, neuronal immaturity, and prolonged hyper-locomotor activity in mice.

### 8. ΔFosB is upregulated by Cyclin B/Cdk1 activity and mediates persistent genomic remodeling

We next explored mechanisms underlying long-term genomic changes by performing motif enrichment analysis^55^ of ATAC-seq data (Fig. 1h, i). We found that open chromatin peaks showed the highest similarity to ChIP-seq profiles of AP-1 transcription factors, dimers composed of Fos and Jun family proteins, such as Fra1, Atf3, Batf, and JunB (Extended Data Fig. 8a), suggesting that AP-1 mediates long-term genomic remodeling. We focused on ΔFosB, a truncated FosB isoform, as a candidate driver of these long-term changes due to its exceptionally long half-life (∼208 hours^56^). We analyzed its expression time course following REPOPS. After 3-day REPOPS, ΔFosB was transiently increased at 24 hours but returned to baseline at 14 and 40 days (Fig. 6a). In contrast, after 10-day REPOPS, ΔFosB was elevated at 24 hours and remained high at 14 and 40 days (Fig. 6a), suggesting that chronic repetition of activation induces long-lasting elevation in ΔFosB. This sustained elevation likely reflects increased baseline expression of Fosb rather than residual REPOPS-induced ΔFosB (Fig. 6a Extended Data Fig. 8b). Notably, ΔFosB levels negatively correlated with calbindin expression (Fig. 6b), which is consistent with previous reports^21^, suggesting that prolonged ΔFosB upregulation contributes to pseudo-immaturity of DG neurons (see also Extended Data Fig. 9).

**Figure 6.**
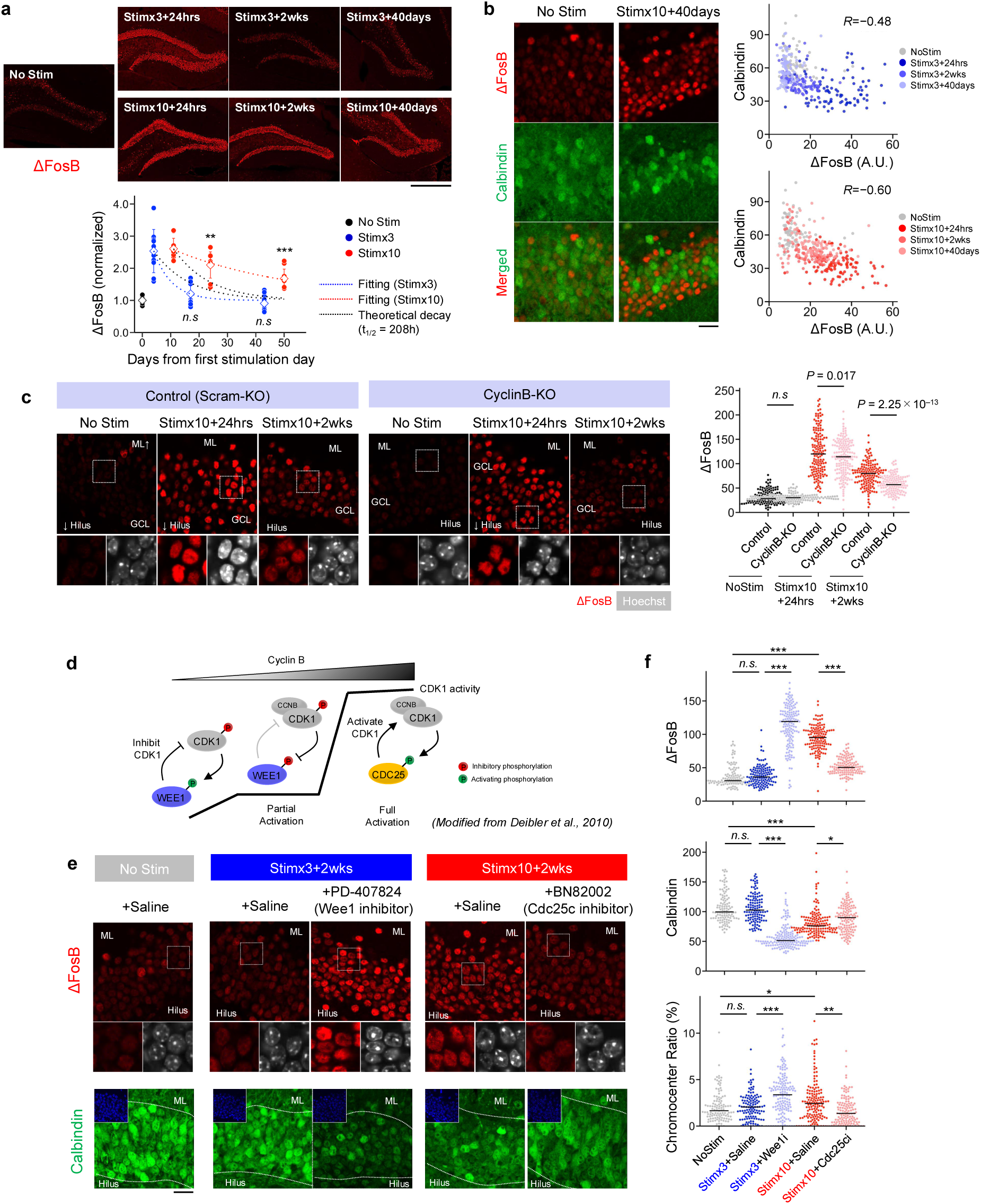
Cyclin B/Cdk1 activity mediates long-term ΔFosB expression and pseudo-immaturity of neurons. **a,** Top, ΔFosB immunostaining in the DG. Scale bar, 500 μm. Bottom, time course of ΔFosB expression. Day 0 denotes the first stimulation day. Values were normalized to the mean of the No Stim group. The black dashed line indicates the theoretical decay of ΔFosB (half-life = 208 h); blue and red curves represent exponential fits for the Stim×3 and Stim×10 groups, respectively. n = 9–11 slices per group. One-sample *t-*tests were performed between observed values and the theoretical decay curve; \*\*\**P* < 0.001. **b,** Immunostaining of ΔFosB and calbindin in the DG (same data as in panel a). Only No Stim and Stim×10+40 days are shown. Scale bar, 20 μm. Right, scatter plot showing the correlation between ΔFosB and calbindin immunoreactivity. **c**, ΔFosB immunostaining in the DG of Scram-KO and CyclinB-KO mice (same mice as in Fig. 5). Bottom panels show enlarged views of the white boxes. Grayscale indicates Hoechst staining. Scale bar, 20 μm. Beeswarm showing ΔFosB immunoreactivity (n = 129–165 cells per group). Black bars indicate the median. One-way ANOVA: *F*_(5, 859)_ = 270.94, *P* < 2.0 × 10⁻¹⁶; post hoc comparisons with Tukey’s test. **d,** Schematic of Cyclin B/Cdk1 activation (modified from Deibler *et al.* 2010^104^). Cdk1 is activated as cyclin B accumulates. This activation is inhibited by Wee1 and promoted by Cdc25c. **e,** Top, ΔFosB immunostaining in the DG. Middle, enlarged view of the boxed region. Grayscale indicates Hoechst staining. Bottom, calbindin immunostaining. Scale bar, 20 μm. **f,** Beeswarm plots showing ΔFosB, calbindin, and chromocenter ratio. n = 114–156 cells per group. Black bars indicate the median. One-way ANOVA: ΔFosB: *F*_(4, 649)_ = 405.45*, P* < 2.0 × 10⁻¹⁶; Calbindin: *F*_(4, 655)_ = 117.65, *P* < 2.0 × 10⁻¹⁶; Chromocenter ratio: *F*_(4, 655)_ = 14.63, *P* < 2.0 × 10⁻¹⁶. Multiple comparisons were corrected using Tukey’s test.

To examine whether cell cycle reentry is involved in this sustained ΔFosB upregulation, we analyzed its expression in CyclinB-KO mice after REPOPS. At 24 hours after 10-day REPOPS, ΔFosB was similarly increased in both CyclinB-KO and Scram-KO groups, indicating that cyclin B is dispensable for acute induction (Fig. 6c). In contrast, at two weeks, ΔFosB was significantly lower in CyclinB-KO than in Scram-KO, suggesting that cyclin B expression is necessary for the sustained elevation of ΔFosB. To further assess the role of Cyclin B/Cdk1, we performed REPOPS while continuously perfusing its modulators*—*PD-407824 (a Wee1 inhibitor) or BN82002 (a Cdc25c inhibitor)*—*via an optofluidic neural probe (Extended Data Fig. 2a) and measured ΔFosB expression (Fig. 6d). Two weeks after 3-day REPOPS, ΔFosB was not elevated, but PD-407824 infusion significantly increased its expression (Fig. 6e). Conversely, after 10-day REPOPS, ΔFosB remained elevated for two weeks, but this increase was suppressed by BN82002 (Fig. 6e). Calbindin showed a complementary pattern: it remained unchanged two weeks after 3-day REPOPS but decreased with PD-407824; whereas it was reduced after 10-day REPOPS but reversed by BN82002 (Fig. 6e). These findings indicate that Cyclin B/Cdk1 activity is essential for sustained elevation of ΔFosB and the associated reduction in calbindin. Thus, repeated neuronal activation and cell cycle reentry induce long-lasting and stable changes in activity-dependent transcription factors, global genomic architecture, and cellular maturity.

## Discussion

In this study, we tested the hypothesis that brain stimulation methods, such as electroconvulsive therapy (ECT) and repetitive transcranial magnetic stimulation (rTMS), may achieve beneficial clinical effects by reprogramming cellular state into an immature but stable configuration. We developed an ECT-like optogenetic stimulation, REPetitive OPtogenetic Stimulation (REPOPS), which reproducibly induces “dematuration” as indexed by global changes in the cellular transcriptome and 3D genome structure. Most of these changes were transient when REPOPS was repeated for three consecutive days, but could persist for more than one month after 10 consecutive days of activation, consistent with the 6–12 stimulus repeat regimen for ECT. RNA-seq data of human DG also showed that ECT induces immature-like gene expression patterns in patients, resembling the effects by REPOPS. Underlining the clinically relevant behavioral effects of REPOPS, we discovered a novel mechanism where our stimulation protocol induces cell cycle re-entry resembling the G_2_/M-phase-like nuclear structure. These changes were reversed by deletion of Cyclin B, a master regulator of the G_2_/M phase transition. We further found that activation of Cyclin B/Cdk1 stabilizes ΔFosB expression, thereby promoting sustained reorganization of genome structure. Our results establish a candidate cellular mechanism underpinning the beneficial clinical effects of ECT and offer a novel cell-based perspective for the development of brain stimulation therapies.

Our findings reveal that optogenetic stimulation of hippocampal granule cells can trigger coordinated changes in gene expression, nuclear structural plasticity, and G_2_/M cell cycle re-entry, which collectively define an immature-like cell state. The evidence supporting G_2_/M cell cycle re-entry includes the increased expression of M phase-related genes such as Cyclin A2, Cyclin B1, Cyclin B2, and Cdc25c, and changes in nuclear structure, including nuclear lamina disruption, an increase in pH3, and chromocenter enlargement. In the literature, cell cycle re-entry has been implicated in several neuropsychiatric and neurological disorders^57,58^. For example, aberrant expression of cell cycle-related genes, such as Cyclin A, Cyclin B, and Cdk1, has been reported in Alzheimer’s disease^57,59–63^ and temporal lobe epilepsy^64^, as well as in animal models of risk events for these diseases, such as epileptic seizures^65,66^, ischemia^67^, and encephalitis^68^. Nuclear alterations related to the cell cycle, such as nuclear lamina disruption and increased pH3, are observed in the brains of patients and animal models of Alzheimer’s disease^58,69,70^ and epileptic seizures^71^. Recent large-scale GWAS studies identified several M phase-related genes (e.g., *CCNB2, CDC25C, CENPM, CENPT, MAD1L1,* and *STAG1*) near the risk loci for schizophrenia^72,73^. While these diseases have diverse etiologies, neural hyperexcitability and cell cycle-like processes may be common contributors to their pathophysiology^74–77^. This study is the first, to our knowledge, to show a causal link between chronic neural activation and G_2_/M cell cycle re-entry. Interestingly, the cell cycle re-entry observed in various diseases is typically associated with apoptosis^78,79^; however, our REPOPS method did not induce significant cell death (Extended Data Fig. 1c). REPOPS triggered nuclear structural dynamics, but they were less pronounced than those in dividing cells during the late M phase^79,80^ and there were no fully compacted chromosomes (Fig. 4e, 4g), suggesting that neurons were arrested in a G_2_ to early M phase-like state for at least two weeks. Importantly, we do not assert that this immature-like cellular state is a perfect regression to developmental immaturity but rather an analog state of immaturity, a state that is phenotypically immature relative to the mature state. Another key finding of this study is the involvement of ΔFosB in neuronal dematuration. ΔFosB has previously been reported to be highly expressed in the striatum of addiction models^81,82^, as well as in the DG of patients with epileptic seizures and Alzheimer’s disease^21,83^. Owing to its long half-life (∼208 hours)^56^, ΔFosB is widely believed to mediate long-lasting cellular changes. However, given this half-life, stimulus-induced ΔFosB would be expected to decay to ∼4% of its peak level within 40 days. Therefore, the marked increase observed in the Stimx10+40day group (Fig. 6a) is unlikely to reflect residual ΔFosB from prior stimulation, but rather a sustained elevation of its basal expression (Extended Data Fig. 8b). This persistent up-regulation of ΔFosB likely contributes to long-lasting 3D genome reorganization and neuronal dematuration. We also suggest that REPOPS triggers repeated calcium influx that initiates dematuration and cell cycle re-entry (Extended Data. Fig. 2a—c). Activation of calcium signalling is essential for cell cycle progression^84,85^; therefore, REPOPS-stimulated neurons may transduce a specific pattern of calcium signaling that causes cell cycle re-entry. Further studies will clarify this signaling pattern, and we predict that variations of REPOPS methodology may enable fine-scale external control of cell dematuration *in vivo*.

ECT is well documented as an effective treatment for psychiatric disorders such as severe depression and anxiety. The current results demonstrate that cellular dematuration triggered by an ECT-like stimulus can produce positive behavioral phenotypes, including decreased depression-like symptoms and increased motor activity, similar to the clinical ECT effects. An important question is how these behaviors are enabled by neuronal stimulation at the neural circuit level. REPOPS increased the number of speed cells and the speed coding accuracy in the DG (Fig. 3i–k). Prior studies indicate that activation of DG neurons can elicit robust exploration of the environment^48^, and our observation of increased speed-related activity in the DG may account for the increased locomotor activity observed in the open field (Fig. 2a, 2b) and home cage (Fig. 2c). Activation of DG neurons is also known to suppress depression-like behaviors^86,87^, therefore, increased speed-related activities of DG neurons may explain the link between ECT-like stimulation and the reduction in depression-like behavior observed in the tail suspension and forced swim tests (Fig. 2d, 2e). In contrast to these beneficial effects, a potential negative side effect of ECT is memory loss, including retrograde and anterograde amnesia^88,89^. Indeed, we observed a deficit in remote memory, or the ability to recall information from neocortex into the hippocampal circuit (Extended Data Fig. 3a and Supplementary results 2). REPOPS caused long-term (> 2 weeks) alterations in synapse-related genes, including *Gria1*, *Gria2*, *Camk2a*, *Camk2b*, and *Arc* (Fig. 1e). Reanalysis of RNA-seq data from human patients with and without ECT also revealed significant changes in the expression of *GRIA1, GRIA2,* and *ARC* (Fig. 1g). These genes play pivotal roles in synaptic and homeostatic plasticity^90–92^ and their decline may be related to memory impairment due to neural hyperexcitation. Long-term down-regulation in the expression of synapse-related genes may interfere with the development of place cells and spatial information coding in the DG (Fig. 3f–h) and CA1^16,93–95^. Such REPOPS-dependent deficit in neural plasticity may share a common mechanism with ECT-induced amnesia. Thus, ECT-like REPOPS alters neural coding, leading to both decreased depressive-like behaviors and memory deficits. Further research will address the ECT stimulus conditions that do not impair memory and learning, but preserve beneficial clinical therapeutic effects.

Our findings also provide a novel perspective on the mechanism of action of antidepressant therapy. In this study, we found that REPOPS-triggered cellular and behavioral effects share intriguing similarities to previously reported behavioral effects and neuronal dematuration induced by ECS or selective serotonin reuptake inhibitors (SSRIs)^9,10,39,40^. Chronic SSRI treatment may reactivate the plastic state in the adult brain, comparable to that observed during development^14,41^. For example, in the adult visual cortex, chronic fluoxetine reopens a critical period of plasticity for the recovery of visual function^96,97^, and, in the hippocampus, increased neural circuit plasticity can promote fear memory extinction^14,98^. SSRIs may regulate neural plasticity by increasing the environmental susceptibility of neural circuits, thereby reducing depression via exposure to favorable environments^99,100^. This form of “meta-plasticity”^101,102^, or the induction of changes in neural plasticity itself, can also manifest in opposite forms by neural hyperactivity. A recent study showed that recurrent seizures cause a long-term reduction in neuronal excitability, indicating decreased neural plasticity^103^. In the current study, REPOPS for 10 days inhibited spatial map formation by exposure to the environment for more than two weeks (Fig. 3f–h), suggesting an altered meta-plasticity for spatial coding. Thus, neural overactivation and dematuration can alter the meta-plasticity of neural circuits, which may underlie the effects of antidepressant treatments such as ECT and rTMS, or the development of neuropsychiatric disorders involving neural hyperexcitability in their etiology. A key insight from the present work is that the immature cellular state appears to be bistable; that is, it can serve as a reversible therapeutic intermediate, as observed in ECT, or consolidate into an irreversible pathological lesion, as seen in disease models.

In summary, we propose a working model for the mechanism of brain stimulation therapies in the treatment of psychiatric disorders. This class of brain diseases may represent neuronal populations trapped in various energy minima of metabolic and/or synaptic inflexibility, such as the loss of plasticity by the pathological closure of neural circuits to environmental stimulation, as discussed above. In this model, repetitive brain stimulation methods like ECT or rTMS trigger patterned calcium influx and signaling to drive cellular dematuration that temporarily resets cellular metabolic and synaptic homeostasis to restore normal neural circuit flexibility and beneficial clinical outcomes like anti-depressive behavior. This immature-like state can persist for more than one month, and excessive neuronal activation may eventually reinforce rather that reset metabolic or synaptic dysfunction in a vicious cycle associated with chronic psychiatric disorders associated with hyperactivity. Therefore, our model suggests that the immature-like cell state may serve as a bistable switch for cellular health at moderate levels of electrical brain stimulation, or disease progression at higher levels. We believe that these findings open a window to novel methods and protocols for artificial neural stimulation that may enable cell reprogramming for brain health.

## Materials and Methods

### Animals

*ROSA26-CAG-stop^flox^-ChR2(H134R)-EYFP* (Strain#012569; RRID:IMSR_JAX:012569) and *POMC-Cre* (Strain #010714; RRID:IMSR_JAX:010714) were obtained from Jackson Laboratories (Bar Harbor, Maine). These mouse lines were crossed to generate POMC-Cre::ChR2-EYFP mice, which express ChR2-EYFP selectively in the granule cells (GCs) of the dentate gyrus (DG) (Fig. 1a). Mice were used for experiments at 3–4 months of age for RNA-seq and ATAC-seq experiments, and at 6–12 months for other experiments. Three or four mice were housed in one cage and switched to individual housing 1–2 weeks prior to the experiment. They were housed with a 12 hr light/dark cycle (lights on at 7:00 a.m., off at 7:00 p.m.) with access to food and water *ad libitum*. The room temperature was maintained at 23 ± 2℃. All the experiments were conducted during the light period. All animal experiments were approved by the Institutional Animal Care and Use Committee of Fujita Health University based on the Law for the Human Treatment and Management of Animals and the Standards Relating to the Care and Management of Laboratory Animals and Relief of Pain. Every effort was made to minimize the number of animals used.

### Stereotactic Surgery

#### Optic fiber implantation

For optogenetic stimulation, a blue-light optic fiber was implanted into the dorsal DG of adult POMC-Cre::ChR2-EYFP mice using conventional methods^105^. Briefly, adult mice were anesthetized with 1.5–3% isoflurane at an oxygen flow rate of 1 L/min. The head fur was shaved and the incision site was sterilized with 70% ethanol before the surgical procedure. The mice were mounted on a stereotaxic device (Narishige, Tokyo, Japan). After the scalp was incised and pulled aside, a 1.0 mm diameter craniotomy was created with a surgical drill on the skull above the implantation site. Using the stereotaxic device, a 250 μm diameter optic fiber (470 nm, fiber diameter 250 μm; TeleC-B-3mm-250μm, Bioresearch Center, Aichi, Japan) was precisely inserted into the target site (2.0 mm posterior to bregma (AP), 1.5 mm lateral to midline (ML), and 1.5 mm ventral to bregma (DV)) and fixed to the skull with dental cement. After implantation, the scalp was sutured and treated with povidone-iodine. To avoid the effects of surgery on mouse behavior, a recovery period of 5–10 days after surgery was provided when performing behavioral experiments.

#### Stereotactic injection of AAV9

For the genome editing experiment (Fig. 5), a mixture of adeno-associated virus (AAV) carrying Cas9 and sgRNA was injected into the DG immediately before optic fiber implantation. For dominant-negative inhibition of ΔFosB experiment (Extended Data Fig. 9), we injected an AAV expressing ΔJunD (a kind gift from A.J. Robison) was injected. Anesthetization and craniotomy were performed in the same way as described above. A glass-capillary pipette (25 μm inner-diameter tip) filled with the virus suspended in mineral oil was inserted into the target site (2.0 mm posterior to bregma (AP), 1.5 mm lateral to midline (ML)) through the surgical drill on the skull. A 200 nL of 1:3 AAV mixture of AAV9-U6-sgRNA (*Ccnb1*-*Ccnb2* or Scramble)-hSyn1-mCherry (2.0 × 10^13^ vector genomes vg/ml) and AAV9-pMecp2-HA-SpCas9 (2.0 × 10^13^ vg/ml) was injected at 200 nL/min using a microinjection pump (Nanoliter 2010, WPI, FL) at three depths (1.5 mm, 2.0 mm, and 2.5 mm). After each injection, a glass pipette was held in place for 1–2 min before retraction to prevent leakage. Shortly after virus injection, the optic fiber was embedded in the same site using the procedure described above.

#### Implantation of injection cannula

In the experiment that combined optogenetic stimulation and drug injection (Fig. 6d–f and Extended Data Fig. 2), a guide cannula (24-gauge, 11 mm length, C316G, Plastics One) was implanted into the dorsal DG using the same procedure as that used for optic fiber implantation.

### Optogenetic Stimulation and drug infusion

For optogenetic stimulation under freely moving conditions, we employed a wireless LED illuminator, Teleopto (BioResearch Center, Nagoya, Japan). A 2.0g infrared receiver (TeleR-2-P, BioResearch Center) was attached to an optic fiber implanted in the mouse head, whereas in the control group, a dummy receiver was used as a substitute. This receiver was signalled by an infrared generator (TeleEmitter-C, BioResearch Center) to light up the optic fiber. The light intensity varied depending on the custom-made fiber, typically within a range of 3–5 mW. We set the stimulation protocol to 10 msec pulses at a frequency of 10 Hz, utilizing a pulse generator (STOmk-2, BioResearch Center). These photostimulation intensities are sufficiently weak to avoid tissue damage^106^.

For optogenetic stimulation with an intracerebral drug infusion (Fig. 6d–f and Extended Data Fig. 2), we employed injection cannulas (31-gauge, 11 mm length, C316I, Plastics One) and customized optic fibers designed to fit the implanted guide cannula (Bio Research Center). For acute injection (Extended Data Fig. 2), the injection cannula was attached to the implanted guide cannula, and 2uL of the drugs was injected over approximately 5 min. The Ca^2+^ blockers consisted of 100 mM NBQX (ab120046, Abcam), 100 mM AP5 (ab120003, Abcam), 100 mM mibefradil (ab120343, Abcam), and 100 mM nimodipine (ab120138, Abcam) diluted in saline (Extended Data Fig. 2c). Kinase inhibitors, K252a (sc-200517, Santa Cruz), SB218078 (sc-203692, Santa Cruz), Dasatinib (sc-218081, Santa Cruz), and Ro-32-0432 (sc-3549, Santa Cruz), were used at the same concentration (1.0 mM). Five minutes after the injection, the injection cannula was gently detached, and a customized optic fiber was inserted through the guide cannula. For chronic drug infusion (Fig. 6d–f), an osmotic pump (Alzet model 1004) was implanted subcutaneously and connected to a guide cannula. The pump was filled with PD-407824 (sc-203669, Santa Cruz) or BN82002 (#217691, Calbiochem) and delivered the drug at a constant rate. For the effective infusion of drugs into brain tissues *in vivo*, the concentration of the solution was set to 100–1000 times higher than that used for bath applications in acute slice cultures according to previous studies^107^. Stimulation was performed in the same manner as described above. All procedures were conducted in an awake state.

### RNA-Sequencing Library Construction, Sequencing, Mapping, and Data Analysis

The dentate gyrus was rapidly dissected from the adult mouse hippocampus under a dissection microscope as previously described^108^. Previous studies have shown that such preparations are highly enriched with mature dentate granule neurons (–90% NeuN^+^ neurons^30,109^; see also Extended Data Fig. 7a). Total RNA from dissected tissue samples was extracted using NucleoSpin RNA (Macherey-Nagel, Duren, Germany), according to the manufacturer’s instructions. The RNA quality was assessed with an Agilent 2200 TapeStation (Agilent Technologies, Santa Clara, CA, United States). Sequencing libraries were prepared using the TruSeq Stranded mRNA Sample Prep Kit (Illumina, San Diego, CA, USA) and sequenced as 100-bp paired-end reads on the Illumina HiSeq 2500. Sequenced data were mapped onto the Mus musculus genome (GRCm38/mm10) using TopHat (v2.0.14). Fragments per kilobase per million (FPKM) values were computed by Genedata Profiler Genome (v10.1.12). Genes with low expression (FPKM < 0.1) were excluded from the analysis to avoid an excessively high FDR due to low expression levels.

### ATAC-Sequencing Library Construction, Sequencing, Mapping, and Data Analysis

The dentate gyrus of adult mice was dissected in the same manner as that used for RNA-seq. The dentate gyrus was slowly frozen in a freezing box (Cat. No. 5100-0001; Mr. Frosty Freezing Container, Thermo Fisher Scientific). Flash-frozen tissue was sent to Active Motif to perform the ATAC-seq assay. The tissue was manually dissociated, isolated nuclei were quantified using a hemocytometer, and 100,000 nuclei were tagmented as previously described^110^, with some modifications based on Corces et al. 2017^111^ using the enzyme and buffer provided in the Nextera Library Prep Kit (Illumina). Tagmented DNA was purified using the MinElute PCR purification kit (Qiagen), amplified with 10 cycles of PCR, and purified using Agencourt AMPure SPRI beads (Beckman Coulter). The resulting material was quantified using the KAPA Library Quantification Kit for Illumina platforms (KAPA Biosystems) and sequenced using PE42 sequencing on a NextSeq 500 sequencer (Illumina). For the analysis of ATAC-seq data, we first aligned reads using the BWA algorithm (mem mode; default settings). Duplicate reads were removed, and only reads mapping as matched pairs and uniquely mapped reads (mapping quality >= 1) were used for further analysis. Alignments were extended in silico at their 3’ends to a length of 200 bp and assigned to 32-nt bins along the genome. The resulting histograms (genomic “signal maps”) were stored in the bigWig files. Peaks were identified using the MACS 2.1.0 algorithm at a cutoff of *P*-value 1.0 × 10^−7^, without the control file, and with the –nomodel option. Peaks on the ENCODE blacklist of known false ChIP-Seq peaks were excluded. Signal maps and peak locations were used as input data for Active Motif’s proprietary analysis program, which creates Excel tables containing detailed information on sample comparisons, peak metrics, peak locations, and gene annotations. For differential analysis, reads were counted in all merged peak regions (using Subread) and the replicates for each condition were compared using DESeq2. IGV was used to visualize the raw intensities^112^. Statistical analyses were performed using an in-house R script, unless otherwise specified.

### Immunohistochemistry

#### Immunostaining of brain sample with PFA fixation

The mice were deeply anesthetized with isoflurane and transcardially perfused with PBS, followed by 4% PFA in PBS. Brains were dissected, immersed overnight in 4% PFA in PBS, and cryoprotected by incubation in 20% sucrose in PBS for three days at 4 ℃. After cryoprotection, the brains were mounted in Tissue-Tek optimal cutting temperature compound (Miles, Elkhart, IN) and frozen in liquid nitrogen. Brain samples were cut into 10-μm-thick coronal sections using a microtome (CM1850; Leica Microsystems, Wetzlar, Germany). The sections were stained with the following antibodies: Calbindin (1:1000, rabbit, polyclonal, Synaptic Systems Cat# 214 002, RRID:AB_2068199), Cyclin B (1:500, mouse, monoclonal, Thermo Fisher Scientific Cat# MA1-155, RRID:AB_2536863), phospho-Histone H3 (1:500, rabbit, polyclonal, Millipore Cat# 06-570, RRID:AB_310177), HA-tag (1:500, rabbit, polyclonal, Cell Signaling Technology Cat# 3724, RRID:AB_1549585), ΔFosB (1:1000, rabbit, monoclonal, Cell Signaling Technology Cat# 14695, RRID:AB_2798577), Calbindin (1:1000, rabbit, polyclonal, Synaptic Systems Cat# 214002, RRID:AB_2068199), NeuN (1:500, mouse, monoclonal, Millipore, Cat# MAB377X, RRID:AB_2149209), GFAP (1:500, rabbit, polyclonal, Sigma-Aldrich, Cat# G9269, RRID:AB_477035), Iba1 (1:500, rabbit, polyclonal, FUJIFILM, Cat# 019-19741, RRID:AB_839504), JunD (1:500, rabbit, polyclonal, Abcam, Cat# ab28837, RRID:AB_2130167), and tri-methyl-Histone H3 (Lys9me3) (1:500, mouse, monoclonal, FUJIFILM Cat# MABI0308). Before the first antibody reaction, antigen retrieval was performed in 120℃ autoclave for 5 minutes in 0.01 M sodium citrate buffer (S1699, Target Retrieval Solution, Dako). Autoclave at 120℃ is essential, especially for detecting weak signals of nuclear antigens (e.g., pH3 and Cyclin B). This procedure also eliminates EYFP/mCherry fluorescence, thus preventing overlap with secondary antibody staining with Alexa 488/594. After antigen retrieval, sections were cooled at room temperature for 2 hours. Sections were pre-incubated for 1 hour at room temperature in 5% skim milk in PBST for blocking, and then incubated overnight at 4℃ in PBS containing the primary antibodies. The next day, secondary antibody reactions were performed for 1 hour at room temperature using Alexa Fluor 488-, Alexa Fluor 594-, and Alexa Fluor 647-conjugated secondary antibodies (Invitrogen). Nuclear staining was performed using Hoechst 33258 (Polysciences, Warrington, PA).

#### Immunostaining of brain sample with acetone fixation

For Lamin B1 staining (1:200, mouse, monoclonal, Thermo Fisher Scientific Cat# MA1-06103, RRID:AB_2281281), a method omitting PFA perfusion was employed^113,114^. In this protocol, brains were perfused with PBS alone, without PFA. Immediately after PBS perfusion, the dissected brains were mounted in Tissue-Tek compound and frozen in liquid nitrogen. Brain samples were cut into 10-μm-thick coronal sections using a microtome. Before the first antibody staining, the brain sections were fixed with ice-cold acetone. Antigen retrieval before the blocking step was omitted and subsequent procedures were performed in the same manner as for immunostaining of the fixed brain samples.

#### TUNEL staining

In Situ Cell Death Detection Kit TMR red (Roche, 12156792910) was used for the detection of apoptotic cells. Brain sections were rinsed in PBS, permeabilized with 0.1 % Tween for 30 min, and incubated with TUNEL reaction reagent (TdT Enzyme and Labeling Safe Buffer) and Hoechst for 60 min at room temperature. Embryo brain tissue was used as the positive control. The localization of TUNEL to the nucleus allows for accurate counting of dead or dying cells.

#### Image acquisition and analysis

Confocal microscopy (LSM 700; Zeiss) was used to obtain images of stained sections. For image analysis, the regions of interest (ROIs) were manually delineated using ZEN (Zeiss) and ImageJ software.

### Behavioral tests

#### Open field test

The open field test was performed in an open field apparatus (40 cm × 40 cm × 30 cm; width, depth, and height, respectively; O’Hara, Japan) made of opaque white plastic, with the center area illuminated to 100 lux by an LED light attached above the ceiling. Before each behavioral experiment, the open field apparatus was cleaned with weakly acidified hypochlorous water (super-hypochlorous water; Shimizu Laboratory Supplies, Kyoto, Japan) to prevent bias due to olfactory cues. All behavioral experiments were conducted in a soundproof room. During each recording session, mouse movements were captured using a CCD camera above the open field arena. Mouse behavior was monitored using a computer screen located outside the room to minimize artifacts caused by the experimenter’s presence. Mouse behavior was recorded at a sampling rate of 10 Hz. Mouse images were automatically processed using the ImageJ plugin (Image OF, freely available on the Mouse Phenotype Database website: http://www.mouse-phenotype.org/software.html) to obtain the location-time sequence for each mouse. From the traces of mouse movements, we calculated the total distance traveled by the mice during the session^115^.

#### Locomotor activity monitoring in home cage

Locomotor activity monitoring was performed on mice that had been operated at least one month before. The system automatically analysed the locomotor activity of the mice in their home cage^116^. The system contains a home cage (29 × 18 × 12 cm) and a filtered cage top, separated by a 13-cm-high metal stand containing an infrared video camera, which is attached to the top of the stand. Each mouse was individually housed in each home cage, and their locomotor activity was monitored 24 hours a day. Outputs from the video cameras were fed into a computer and images from each cage were captured at a rate of one frame per second. The distance traveled was calculated automatically using the ImageJ plugin^115^.

#### Tail suspension test

The tail suspension test was performed to assess depression-related behavior in mice. Each mouse was suspended 30 cm above the floor by the tail in a white plastic chamber (44 cm height × 49 cm length × 32 cm width; O’Hara, Japan) with a video camera mounted on the wall (O’Hara, Japan). The behavior was recorded for 10 minutes. Images were captured at two frames per second using a video camera and transferred to a computer. The area (pixels) within which the mouse moved was measured for each pair of successive frames. When the area was below a certain threshold, the mouse behavior was judged as “immobile.” When the area equaled or exceeded the threshold, the mouse was considered “moving.” The optimal threshold for this judgment was determined by adjusting it to the amount of immobility measured by a trained human observer. Immobility lasting for < 2 seconds was not included in the analysis. Data acquisition and analysis were performed automatically using the ImageJ plugin software^115^.

#### Porsolt forced swim test

The Porsolt forced swim test was performed to assess depression-related behavior. A Plexiglas cylinder (20 cm height × 10 cm diameter) was placed in a test chamber (49 cm height × 44 cm length × 32 cm width; O’Hara, Japan). A video camera was mounted on the ceiling of the test chamber and positioned directly above the cylinder. The mice were placed into a cylinder, which was filled with water (approximately 23°C) to a height of 7.5 cm.

The immobility time was recorded for 10 minutes. Images were captured at two frames per second through the video camera. The immobility status of each mouse was determined in the same manner as for the tail suspension test. Data acquisition and analysis were performed using the ImageJ plugin^115^.

#### Contextual and cued fear-conditioning tests

Fear conditioning tests were conducted to assess contextual memory (Extended Data Fig. 3a). In the conditioning session, each mouse was placed in an acrylic chamber consisting of white (side) and transparent (front, rear, and top) walls (33 × 25 × 28 cm) with a stainless-steel grid floor (0.2 cm diameter, spaced 0.5 cm apart; O’Hara, Japan). The mice were allowed to explore the chamber freely for 120 seconds, and 55 dB white noise then served as the conditioned stimulus (CS) for 30 seconds. During the last 2 seconds of CS presentation, a mild foot shock (0.3 mA, 2 seconds) was delivered as the unconditioned stimulus (US). The mice were subjected to two more CS-US pairings with a 2-minutes interstimulus interval. Approximately 24 hours or one month after conditioning, a context test was conducted for 300 seconds. In the context test, mice were placed in the same chamber in which they had been conditioned. After the context test, a cued test with an altered context was performed for 360 seconds. In the cued test, the mice were placed in a triangular chamber (33 × 29 × 32 cm) made of white plastic walls and floor, which was located in a different sound-attenuating room and allowed to explore the triangular chamber for 180 seconds. Then, the CS was presented during the last 180 seconds of the cued test. In each session, the percentage of freezing and distance traveled was calculated automatically using the ImageJ plugin^115^.

#### Three chamber social interaction test

The testing apparatus consisted of a rectangular three-chambered box and lid with a video camera (O’Hara, Japan; Extended Data Fig. 3b). The dividing walls of the chamber were made of transparent plastic, with small square openings (5 × 3 cm) allowing access to each chamber (20 × 40 × 47 cm). A small round wire cage (9 cm in diameter, 11 cm in height, with vertical bars 0.5 cm apart) was located in the corner of chambers. The test mice were first placed in the middle chamber and allowed to explore the entire test chamber for 10 minutes. Immediately after the 10-minute period, the test mice were placed in a clean holding cage, and a male C57BL/6J mouse (stranger mouse) with no prior contact with the test mice was enclosed in one of the wire cages. The test mice were returned to the middle chamber and allowed to explore for 10 minutes. The time spent in each chamber and the time spent around each cage was automatically measured using ImageCSI software^115^.

### in vivo Ca^2+^ imaging with optogenetic stimulation

#### Viral constructs

Purified and concentrated adeno-associated virus serotype 5 coding GCaMP6f under the synapsin promoter (AAV5-syn-GCaMP6f; #100837-AAV5) and adeno-associated virus serotype 5 coding for ChrimsonR-tdTomato under the synapsin promoter (AAV5-syn-ChrimsonR-tdT) were obtained from Addgene and UNC Vector Core Facility.

#### Viral delivery of Ca^2+^ sensor and opsin

For delivery of a fluorescent Ca^2+^ sensor and red-shifted opsin into dDG neurons, AAV5-syn-GCaMP6f and AAV5-syn-ChrimsonR-tdT were injected into the dorsal DG in adult mice (left hemisphere; 2.0 mm posterior to bregma [AP], 1.0–1.45 mm lateral to midline [ML], and 2.0 mm ventral to bregma [DV]). Briefly, adult mice (> 8 weeks old) were anesthetized with 1.5 to 3% isoflurane at an oxygen flow rate of 1 L/min. The head fur was shaved and the incision site was sterilized with 70% ethanol prior to the surgical procedure. Mice were mounted on a stereotaxic device (catalog no. 51730D, Stoelting), and heat pad (catalog no. BWT-100A, BRC) was placed underneath each mouse to maintain body temperature at 37 °C. After the scalp was incised and pulled aside, a 2-mm-diameter craniotomy was created with a surgical drill on the skull above the injection site. Through a glass-capillary injection pipette (25-µm inner-diameter tip), a mixture of 300nl of AAV5-Syn-GCaMP6f-WPRE-SV40 (titer 4.3 × 10^12^ GC/mL) and AAV5-syn-ChrimsonR-tdT (titer 1.53 × 10^12^ GC/mL) was injected using a microinjection pump (Nanoliter 2010, WPI). After the injection, the scalp was sutured and treated with povidone–iodine. The expression of GCaMP6f is driven by the Syn promoter and is expected to be induced not only in GCs but also in other cell types. We previously reported that the majority (approximately 90%) of neurons expressing GCaMP in the DG are GCs^16^.

#### GRIN lens implantation

At least two weeks after viral injection, a gradient refractive index (GRIN) lens (Inscopix catalog no.1050-004623, 1 mm diameter) was implanted at the same position where the virus was injected. The mouse was mounted on a stereotaxic device and the scalp was removed. After exposing the skull and removing the overlying connective tissue, we made a cranial hole slightly larger than the diameter of the GRIN lens. To secure for an entry path for the GRIN lens, the cortex and white matter were aspirated using a 26-gauge blunt needle attached to a vacuum pump; to prevent aspiration of the GCs, the depth of needle tip depth was set at 1.6 mm from the bregma and the tissue below was carefully aspirated. Saline was continuously applied during aspiration to prevent the tissue from drying. During implantation, the lenses were held in place and slowly lowered using a ProView Implant kit (nVoke, Inscopix, Palo Alto, California, USA) attached to the stereotaxic arm. The GRIN lens connected to the manual stereotaxic arm was slowly lowered (5–10 μm/sec) until the field of view was in focus, referring to the fluorescence intensity of the GCaMP. After temporarily immobilizing the lens with ultraviolet-curable resin (Primefil, Tokuyama Dental), it was fixed to the exposed skull using dental cement (Super-Bond C&B, Sun Medical). The exposed tip of the GRIN lens was covered with dental silicone (Dent Silicone-V, Shofu). After lens implantation, analgesic and anti-inflammatory agents (Flunixin, 2 mg/kg; Fujita) and antibiotics (Tribrissen, 0.12 mL/kg; Kyoritsu Seiyaku) were injected intraperitoneally.

#### Baseplate attachment

At least two weeks after GRIN lens implantation, the baseplate of a miniature microendoscope (nVoke, Inscopix) was attached to the GRIN lens using a previously described method^16^. Briefly, the mice were anesthetized and mounted onto a stereotaxic device, and a baseplate attached to the miniature microscope was placed on the GRIN lens using a gripper (nVoke accessory, Inscopix). The optimal location was determined by monitoring the fluorescent images of GCaMP-expressing neurons (where the largest number of neurons was in focus), and the baseplate was fixed with ultraviolet-curable resin and dental cement at this position. In cases where we failed to identify neurons at this stage (usually due to the failure of GRIN-lens implantation), the baseplate was not attached, and such mice were excluded from the experiment. After the baseplate attachment, a baseplate cover was placed on the baseplate until Ca^2+^ imaging was performed.

#### Ca^2+^ imaging and optogenetic stimulation in freely moving mice

Ca^2+^ imaging of the DG neurons was performed while the mice traveled freely in the OF. Prior to Ca^2+^ imaging, the mice were lightly anesthetized and a miniature microscope (nVoke, Inscopix) was mounted onto the baseplate of the mice. The mice were then placed back in their home cages and transferred to a sound-proof behavioral experiment room. At least 30 min after recovery from anesthesia, the mice were subjected to OF. The Ca^2+^ signals of their DG neurons were obtained at a 10-Hz sampling rate with a 1,440×l,080-pixel resolution. Recordings per day totaled 50 min, consisting of before (0–30 min), during (30–35 min), and after (35–50 min) optogenetic stimulation; no stimulation was introduced in the control group. These 50-min recordings were performed daily throughout the experimental period, and the first 30 min recordings were used for data analysis. The experimental period consisted of before stimulation (Day 1), during stimulation (Day 2-11), and two weeks after stimulation (Day 24-26). We applied EX-LED (filtered with a 435–460 nm excitation filter) for the excitation of GCaMP6f fluorescence (0.4 mW/mm^2^ at the bottom of the GRIN lens) and OG-LED (filtered with a 590– 650 nm excitation filter) for the excitation of ChrimsonR (10 mW/mm^2^ at the bottom of the GRIN lens). A previous study reported that EX-LED at 2.0 mW/mm^2^ resulted in small current changes of approximately 10 pA via activation of ChrimsonR under slice preparation (approximately 130 pA by OG-LED irradiation)^117^. Given that EX-LED used in this study are much lower than these conditions (–0.4 mW/mm^2^) and that under *in vivo* imaging conditions the EX-LED are scattered in the brain tissue and the current changes are much smaller than in slice preparation^117^, unintended activation of ChrimsonR by EX-LED would be negligible.

#### Detection of Ca^2+^ transients

To extract the activity patterns of individual DG neurons from the obtained fluorescent images, we used Inscopix Data Processing Software (IDPS 1.8.0). Briefly, the data files of the raw sequential fluorescent images obtained by nVoke (Inscopix) were imported to the IDPS, and movie trimming, data preprocessing, spatial filtering, and motion correction were applied. The instantaneous fluorescence of the motion-corrected images was then normalized by its average fluorescence over the entire recording period, producing fluorescence-change ratio (Δ*F*/*F*) images. Individual DG neurons were identified using the automated principal component analysis (PCA) and independent component analysis (ICA) segmentation of activity traces. For PCA/ICA segmentation, default parameters, whose values were confirmed to work well across most of the neuronal activity patterns in the cortex and CA1 of the hippocampus, were used (number of independent components, 120; number of principal components, 150; ICA max iterations, 100; ICA random seed, 0; ICA convergence threshold, 0.00001; block size, 1,000; ICA unmixing dimension, spatial; ICA temporal weights, 0.00; and arbitrary units within the IDPS). Finally, the temporal pattern of each timing of DG neuron activity was identified by an event-detection algorithm available in IDPS (event threshold factor, four median absolute deviations; event shortest decay time, 0.2 s), and the temporal pattern of activity was expressed as a time series of binarized signals for further analysis.

### Information coding in individual neurons

#### Definitions of the position and speed of mice

i. Position: The OF arena (which had an area of 40 cm × 40 cm) was represented as 200 × 200-pixel grid. The position of the mouse was determined from the centroid of its shadow on the CCD camera, above the OF. We then assigned a label corresponding to the discrete location of the mouse (e.g., [10, 100]) to each time bin (=1/10sec).
ii. Speed: From the distance traveled between 1 s before and after a given time point, we calculated the speed of the mouse at that moment and assigned this speed (cm/s) to each time bin.

#### Statistical analysis of spatial and speed information

To quantify the tuning specificities of neurons with respect to the position and speed of the mouse, we measured their specificity in terms of the information rate of cell activity, and defined them as (i) spatial and (ii) speed information^16,118^. The Ca^2+^ event rate in Ca^2+^ imaging was considerably lower than that in electrophysiological recordings (average, approximately 0.05 Hz). If the discretization of position and speed is too fine for the number of events in the recorded cells, a proper null distribution cannot be obtained when creating shuffled data for that cell. Therefore, we set the resolution of the discretization of the position and speed to be lower than those that are commonly used.

i. We used a 2×2 square grid to measure spatial information and computed the amount of Shannon information conveyed by a single Ca^2+^ transient about the animal’s position. The spatial information *I* (bits per Ca^2+^ transient) of a cell was calculated as the mutual information score between the occurrence of a single Ca^2+^ transient in the cell and the animal’s behavioral state of position using the following formula:

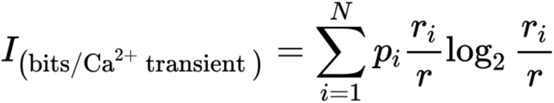 where *i* is the bin number corresponding to the physical parameter (in this case, spatial position in the OF; i = 1–4 from the 2×2 square grid), *N* is the total number of bins, *p* is the probability that the mouse occupied bin *i*, *r_i_* is the mean transient rate at bin *i,* and *r* is the overall mean transient rate.
ii. To measure speed information, we applied the same formula to the speed after discretizing it to a binary state: Run (speed > 1 cm/sec) and Stop (speed < 1 cm/sec).

#### Definitions of place cells and speed cells

To evaluate the statistical significance of the information encoded in individual neurons, we employed the random permutation method as previously outlined^16^. The calcium event data were divided into 1000 segments along the time axis and randomly shuffled to generate permuted calcium event data. This method disrupts the temporal structures of neural activity and the temporal correlations between neural activity and behavioral variables. Using this randomly permuted data, we derived a null distribution for the position and speed information of neurons. Neurons with original information content exceeding the top 95% and 99% of these null distributions were classified as place or speed cells, respectively, indicating cells with significant spatial or speed information^119,120^.

### Decoding position and speed in the open field

To determine how OF behavioral parameters are encoded in the population activities of DG neurons, we trained decoders using machine learning methods to predict the position and speed of mice from the population Ca^2+^ activity. We assigned labels to the discretized behavioral parameters of the mouse (position (cm) and speed (cm/s)) and the binary values of the Ca^2+^ signal (0 or 1) of all neurons in each time bin. We then divided the Ca^2+^ imaging data and behavioral data of the first 30 min (before 5-min of optogenetic stimulation each day) into the first 15 min and last 15 min halves, which were designated as training and test data, respectively. For each pair of behavioral parameters (position and speed) and the value of the Ca^2+^ signal in the training data, we trained the decoders using a machine learning method (the code was obtained and modified from Glaser *et al*., eNeuro, 2020^121^). The decoding accuracy of the position was reported as the mean absolute error of the distance between the predicted and actual positions. In the case of speed, decoding accuracy was reported as the correlation between the predicted and actual instantaneous speeds. This is because speed, unlike position, is not limited to a certain range and the mean absolute error of speed depends on the average locomotion speed of each mouse, which is inappropriate for the evaluation of the decoding error. If the number of neurons recorded in each session was less than 10, the session was excluded from the decoding analysis.

### DNA Constructs and AAV Vectors for in vivo genome editing

For SpCas9 target selection and single-guide RNA (sgRNA) generation, 20-nt target sequences were selected to precede a 5’-NGG protospacer-adjacent motif (PAM) sequence. To minimize off-target effects, CRISPR design tool was used (https://crispr.dbcls.jp/). The backbone plasmid for sgRNA (PX552; pAAV-U6sgRNA(SapI)_hSyn-GFP-KASH-bGH) was obtained from Addgene. *Ccnb*-sgRNA (*Ccnb1* and *Ccnb2* were tandemly connected; Fig. 5b) and *Scramble*-sgRNA nucleotides were synthesized and cloned into AAV backbone plasmids (Vigene Biosciences, Inc, USA). To avoid fluorescence overlap with ChR2-EYFP, the GFP cassette was replaced with mCherry (Fig. 5b). The plasmid for SpCas9 was obtained from Addgene (PX551; pAAV-pMecp2-SpCas9-spA). Concentrated AAV vectors were produced from plasmids for *Ccnb*-sgRNA, *Scramble*-gRNA, and SpCas9, with support from the Gunma University Initiative for Advanced Research Viral Vector Core.

### Stimulated Emission Depletion Microscopy (STED) observation

Brain slices were double-immunostained with anti-tri-methyl-Histone H3 (Lys9me3) antibody (1:500, mouse, monoclonal, MAI0308, FUJIFILM) and anti-phospho-Histone H3 antibody (1:500, rabbit, polyclonal #06-570, Cell Signaling) as the primary antibodies, followed by Alexa Fluor 488- and Alexa Fluor 555-conjugated antibodies (Molecular Probes) as the secondary antibodies. STED images were acquired using a TCS SP8 STED 3X system (Leica Microsystems, Wetzlar, Germany) with 2ch HyDSMD detectors^122^. A white laser (for excitation) and a 660 nm laser with a donut beam (for stimulated emission depletion) were used for STED imaging. For the detection of signals of tri-methyl-Histone H3 (Lys9me3) and phospho-Histone H3, we used excitation wavelengths of 488 and 561 nm, respectively. Raw STED images (pixel size:20 nm) were recorded with a 93 × glycerol immersion objective (Leica HC PL APO 93 × /1.30 GLYC motC STED W) and deconvoluted using the Huygens software (SVI, Hilversum, Netherlands). Image analysis was performed with a custom-developed code using the image processing toolbox in MATLAB (MathWorks, R2022a).

### Data Collection, Pre-processing, and Meta-analysis

All microarray and RNA-seq data used for meta-analysis in this study were obtained from the BaseSpace Correlation Engine (https://japan.ussc.informatics.illumina.com/c/nextbio.nb; Illumina, Cupertino, CA), a publicly available database containing over 100,000 microarray and RNA-seq datasets. The details of all datasets are described in Supplementary Table1. The datasets registered in BaseSpace underwent several preprocessing, quality control, and organization stages. Quality control ensures the integrity of samples and datasets. Genes with a *P* value < 0.05 (without correction for multiple testing) were included in the differentially expressed gene datasets. This sensitivity threshold is typically the lowest used with commercial microarray platforms and the default criterion in BaseSpace analyses^44^. Correction for multiple testing was omitted to minimize false negatives at this stage. Expression values were used to calculate fold changes and *P* values between the two conditions (infants–adults and treated– untreated). To determine fold changes, we divided the expression values of the probes/genes in the test datasets by those in the control datasets. If the fold change was < 1.0, these values were converted to a negative reciprocal or −1/(fold change). Genes with a *t*-test *P* value < 0.05 were imported into the BaseSpace Correlation Engine according to the instructions provided by the manufacturer. The rank order of these genes was determined based on their absolute fold-changes. All meta-statistical analyses comparing two datasets were performed in BaseSpace, and the similarities between any two datasets were evaluated as overlap *P* values using the Running Fisher algorithm^44^.

### Statistical Analyses

Mice of the same age and sex were raised under identical conditions and were randomly allocated to different experimental groups. Blinding was not applied in this study because the animals needed to be controlled by the stimulation conditions. The sample sizes were determined based on those reported in similar publications^30,31^. Welch’s t-test was used to compare the averages. An ANOVA was performed when more than two groups were compared, followed by the Bonferroni post-hoc method. For correlation analysis, we used Pearson correlation analysis for normally distributed data. When pooling the data of neurons from multiple mice per group, a Linear Mixed-Effects Regression model was applied to account for individual differences per mouse. All results of the statistical analyses are summarized in Supplementary Table 2.

## Code and Data Availability

RNA-seq and ATAC-seq data used in this study were deposited in the GEO database with accession codes GSE227200 (RNAseq) and GSE227201 (ATAC-seq), respectively. The RNA-seq data for the post-mortem DG of human patient was obtained from SRP241159. The personal information regarding ECT history was provided by Astellas Pharma Inc. The raw imaging data were uploaded to the Systems Science Biological Dynamics repository (https://ssbd.riken.jp/repository/335). The confocal images used in the analysis were uploaded to a figshare with DOI (https://doi.org/10.6084/m9.figshare.28853303). The codes and processed data supporting the findings of this study are available at https://github.com/tmurano. Additional information is available from the corresponding authors upon request.

## Acknowledgments

We thank the members of the Miyakawa laboratory—Wakako Hasegawa, Chikako Ozeki, Tamaki Murakami, Yoko Kagami and Harumi Mitsuya—for technical assistance, and Hirotaka Shoji for critical input on the design of the Ca²⁺ imaging experiment. We are grateful to Akihiro Yamanaka (Nagoya University and the Chinese Institute for Brain Research) for technical advice on optogenetics. We thank A. Konno, H. Hirai, and the Viral Vector Core at the Gunma University Initiative for Advanced Research for producing AAV9 viral particles. We also acknowledge D. Weinberger and T. Hyde for providing the human DG RNA-seq data. We thank Dr. Charles Yokoyama for his valuable comments, discussions, and editing and for greatly improving our manuscript. This research was supported by the Program for Brain Mapping by Integrated Neurotechnologies for Disease Studies (Brain/MINDS) from Japan Agency for Medical Research and Development, AMED under Grant Number JP21dm0207111, Japan Society for the Promotion of Science (JSPS) Grant-in-Aid for Scientific Research on Innovative Areas (grant nos. JP16H06462), JSPS KAKENHI (grant nos. JP20H00522 and JP22K15204), and the Japan Agency for Medical Research and Development Strategic Research Program for Brain Science Grant JP18dm0107101.

## Author Contribution Statement

T. Murano, K. Takao, and T. Miyakawa designed the experiments and analyses. T. Murano performed RNA-seq, ATAC-seq, histological analysis, in vivo genome editing experiments, and data analyses. T. Murano and K. Takao performed the behavioral experiments. T. Murano and Y. Takamiya performed the Ca^2+^ imaging experiments. M. Namihira and K. Katoh performed the STED microscopy observations. K. Tajinda and M. Matsumoto provided RNA-seq data of post-mortem DG of human patients. T. Murano prepared all the figures and wrote the manuscript. T. Miyakawa supervised all aspects of the study. All authors reviewed the manuscript.

