## Supplementary Figures for "Repetitive Neuronal Activation Regulates Cellular Maturation State via Nuclear Reprogramming"

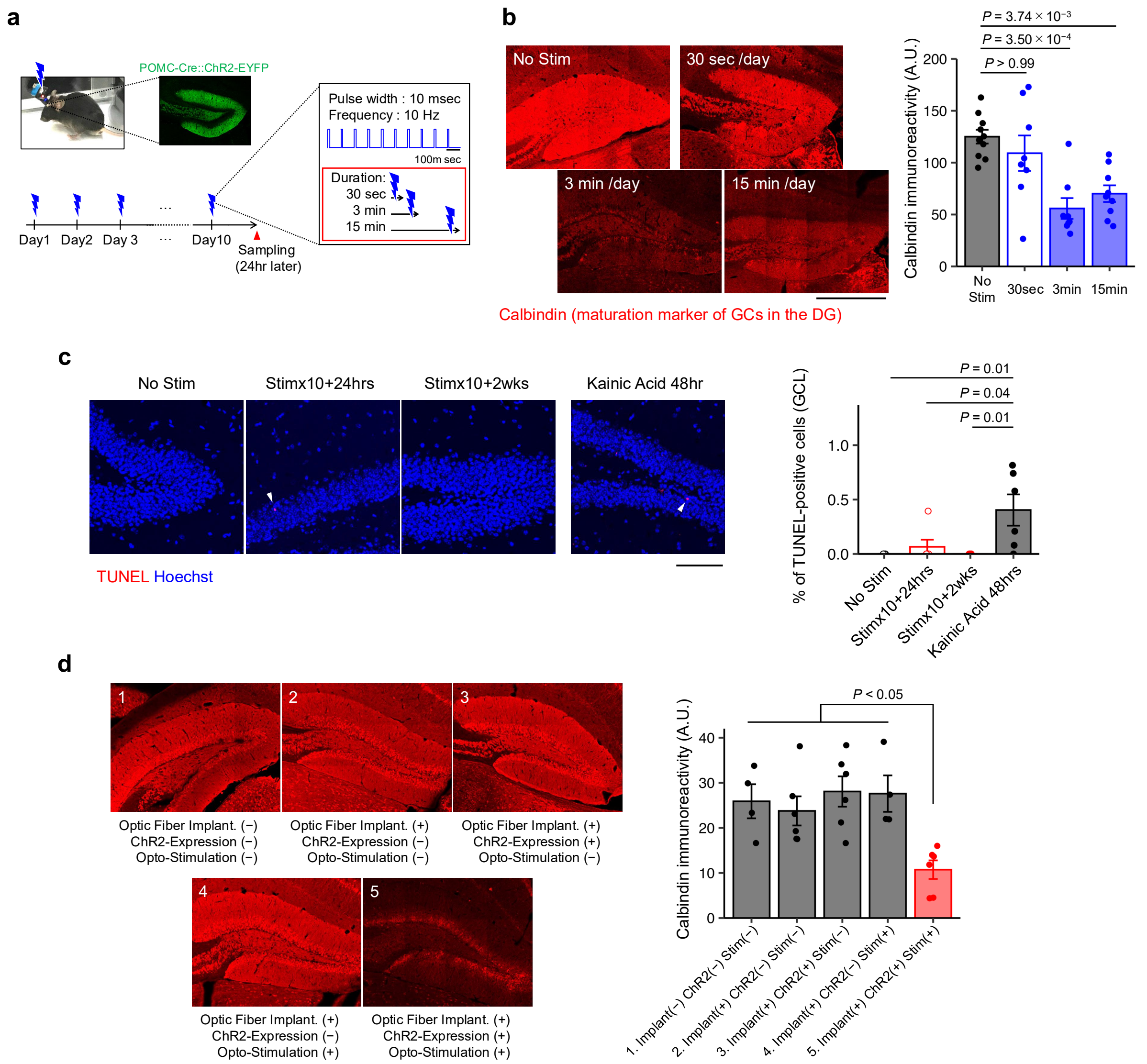


***Supplementary Figure 1. REPOPS induces neural activation without significant cell death***

**a,** REPetitive OPtogenetic Stimulation (REPOPS). ChR2 is exclusively expressed in granule cells (GCs) of the dentate gyrus (DG) under control of the POMC promoter. REPOPS enables frequency- and duration-controlled activation of GCs.

**b,** Calbindin staining (red) in the DG, 24 hours after 10-day REPOPS at various stimulation durations (30 sec, 3 min, 15 min). Scale bar, 500 μm. Bar graph indicates calbindin immunoreactivity. Mean ± s.e.m. (8–10 sections from 4 mice/group). One-way ANOVA, $F_{(3,31)} = 9.75$, $P = 1.35 \times 10^{-4}$; Bonferroni correction was applied for multiple comparisons. Calbindin expression decreased after 3- or 15-minute duration stimulation but not after 30-sec, suggesting that prolonged neural activation is crucial for dematuration.

**c,** TdT-mediated dUTP Nick-End Labelling (TUNEL) staining (red) in the DG of mice treated with 10-day REPOPS or kainic acid (20 mg/kg, 48 hours after intraperitoneal injection).

Arrowheads: TUNEL-positive cells. Scale bar, 500 µm. Bar graph shows % of TUNEL-positive cells in the GCL. Mean ± s.e.m. of 6 mice/group. One-way ANOVA; $F_{(3,20)} = 6.00$, $P = 4.37 \times 10^{-3}$; Bonferroni correction was applied for multiple comparisons.

**d,** Calbindin staining in the DG under five experimental conditions (1−5), representing combinations of optic fiber implantation, ChR2 expression in GCs, and optogenetic stimulation. The bar graph shows calbindin immunoreactivity. Mean ± s.e.m. (4–6 sections from 4 mice/group). One-way ANOVA; $F_{(4,26)} = 0.3244$, $P = 0.858$; Tukey's HSD post hoc test was applied for multiple comparisons.

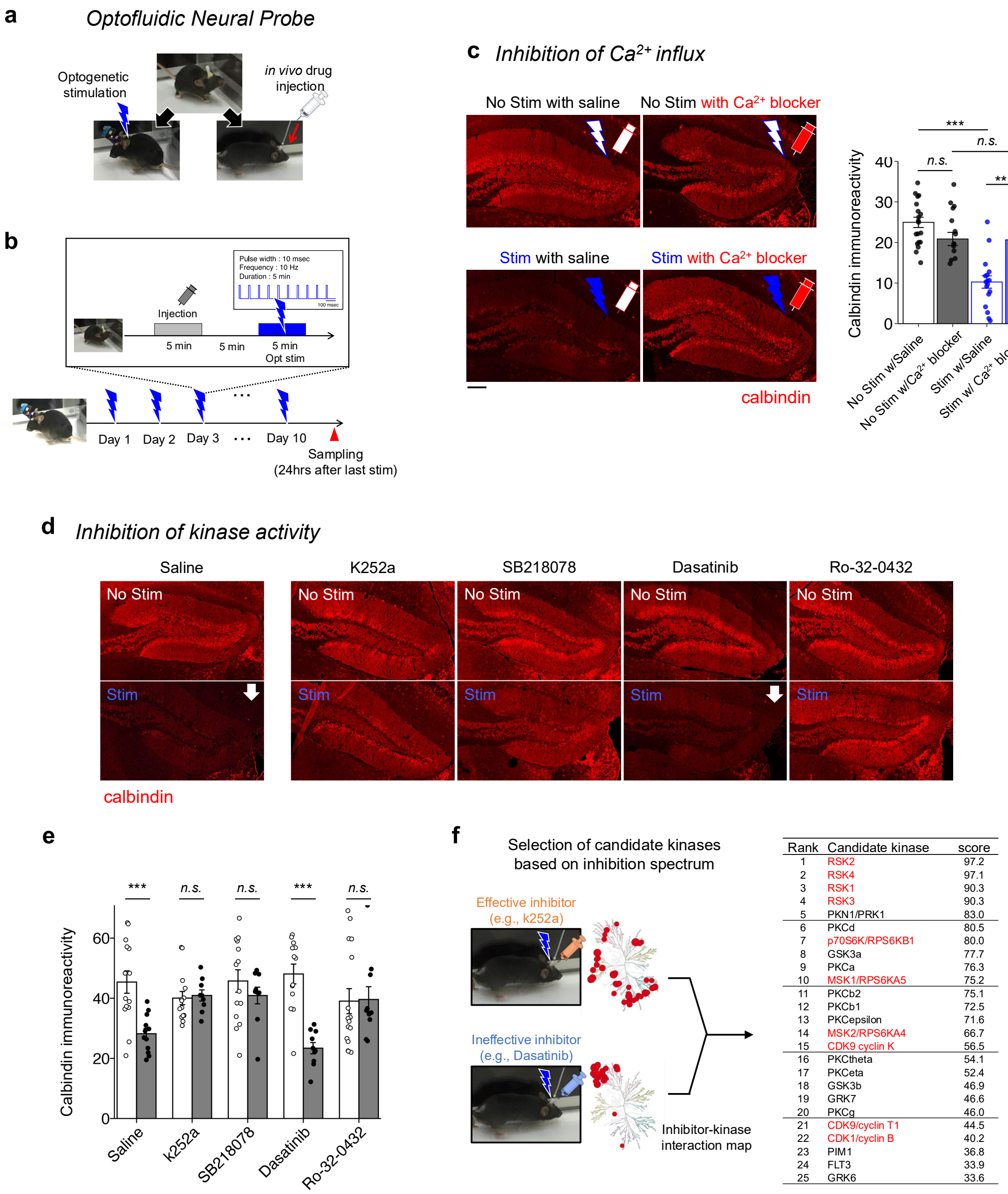


***Supplementary Figure 2. Essential role of $Ca^{2+}$ and kinase activity in REPOPS-induced dematuration***

**a,** Optofluidic neural probe, allowing localized drug injection and optogenetics stimulation through a single cannula implanted in the brain.

**b,** Experimental design. Intracerebral drug infusion into the DG was followed by optogenetic stimulation (5 min/day). This procedure was repeated for 10 consecutive days, and brains were sampled 24 hours after the last stimulation.

**c,** Effect of $Ca^{2+}$ blockers on calbindin expression. $Ca^{2+}$ blockers consisted of NBQX (AMPA-R antagonist), AP5 (NMDA-R antagonist), mibefradil (non-specific blocker of voltage-gated

$Ca^{2+}$ channels), and nimodipine ($Ca_v1$ blocker). Left; immunostaining of calbindin (red); scale bar, 500 μm. Right: bar graph shows calbindin immunoreactivity. Mean ± s.e.m. (14–19 sections from 4 mice/group). One-way ANOVA; $F_{(3,61)} = 19.17$, $P = 7.13 \times 10^{-9}$; Bonferroni correction was applied for multiple comparisons. $^{*}P < 0.05$, $^{**}P < 0.01$, $^{***}P < 0.001$.

**d,** Effect of kinase inhibitors on calbindin expression. Four kinase inhibitors with distinct inhibition spectra (K252a, SB218078, Dasatinib, Ro-32-0432) and saline were injected before optogenetic stimulation. Representative calbindin immunostaining images (red) are shown.

**e,** Bar graph of calbindin immunoreactivity shown in **d.** Mean ± s.e.m. (9–16 sections from 4 mice/group). Two-way ANOVA; $F_{(4,112)} = 5.60$, $P = 3.8 \times 10^{-4}$. Tukey's HSD post hoc test was applied for multiple comparisons.

**f,** Strategy to identify candidate kinases mediating REPOPS-induced dematuration. Left, schematic kinase dendrogram (modified from Anastassiadis et al., 2011[1]) showing the inhibition profiles of the tested compounds; circle size indicates inhibition strength. Right, candidate kinases identified by comparing inhibition spectra of compounds that blocked (K252a, SB218078, and Ro-32-0432) or did not block (Dasatinib) REPOPS-induced effects.

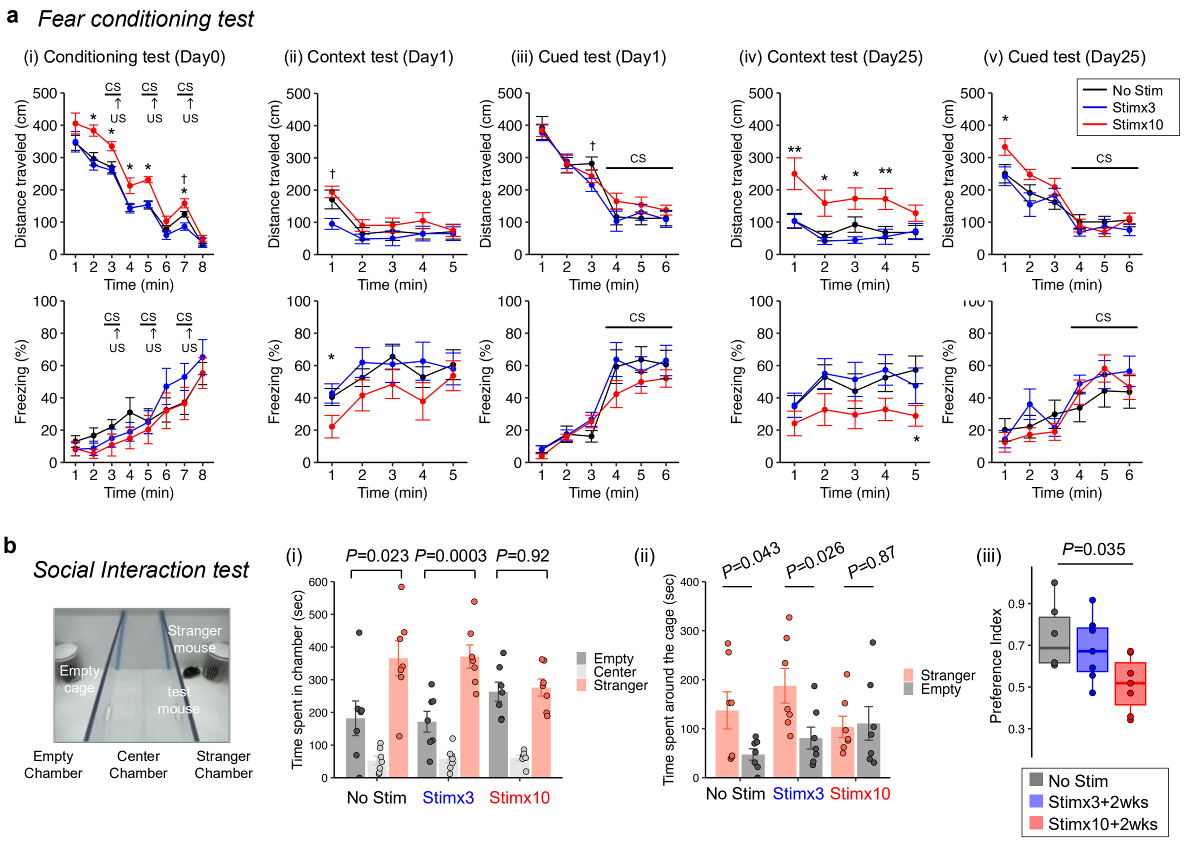


***Supplementary Figure 3. Impaired memory and social behavior following repeated optogenetic stimulation***

**a,** Fear conditioning test. Distance travelled and freezing behavior were assessed during (i) conditioning (Day 0), (ii) context test (Day 1), (iii) cued test (Day 1), (iv) context test (Day 25), and (v) cued tests (Day 25). Data are shown as mean ± s.e.m. for No Stim (n = 10), Stim×3 (n = 8), and Stim×10 (n = 10) groups. Two-way repeated-measures ANOVA followed by Bonferroni correction for multiple comparisons. Asterisks indicate Stim×10 vs. No Stim: *$P < 0.05$, **$P < 0.01$. Daggers indicate Stim×3 vs. No Stim: †$P < 0.05$.

**b,** Social interaction test. (i) Time spent in each chamber (empty, center, stranger mouse) (n = 7 mice/group). One-way ANOVA: No Stim, $F_{(2,18)} = 12.7$, $P = 3.63 \times 10^{-4}$; Stim×3, $F_{(2,18)} = 30.42$, $P = 1.69 \times 10^{-6}$; Stim×10, $F_{(2,18)} = 27.33$, $P = 3.51 \times 10^{-6}$. $P$-values in the graphs were Bonferroni corrected for multiple comparisons. (ii) Time spent around the empty cage vs. the cage with a stranger mouse. Student $t$-test: No Stim, $t_{(12)} = 2.27$, $P = 0.043$; Stim×3, $t_{(12)} = 2.54$, $P = 0.026$; Stim×10, $t_{(12)} = 0.17$, $P = 0.87$. (iii) Preference index, calculated as time spent near the stranger mouse divided by total time spent near both cages. One-way ANOVA, $F_{(2,17)} = 4.18$, $P = 0.0333$. $P$-values were Bonferroni-corrected for multiple comparisons.

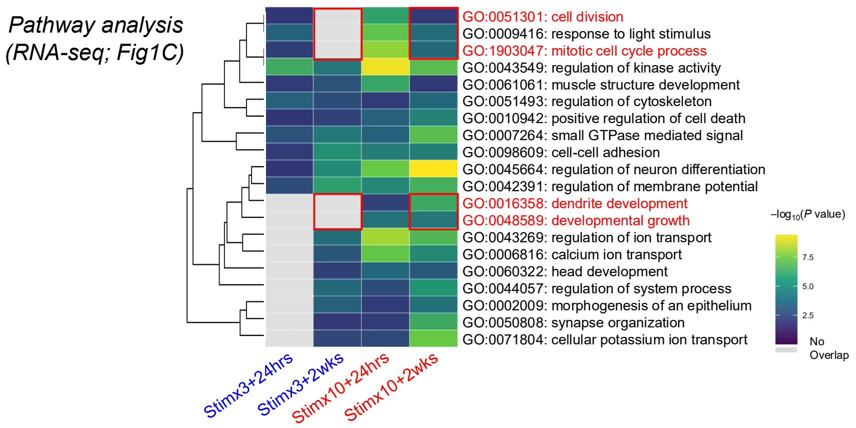


***Supplementary Figure 4. Group analysis of altered gene expression following REPOPS***

Gene Ontology (GO) analysis of RNA-seq data from the optogenetically-stimulated DG (Fig. 1c) using Metascape[2]. Top 20 enriched biogroups are shown. Five biogroups were changed in the Stim×10+2wks group but not in Stim×3+2wks group: Two of them were cell-cycle related (*"cell division"* and *"mitotic cell cycle process"*) and the other two were development-related (*"dendrite development"* and *"developmental growth"*).

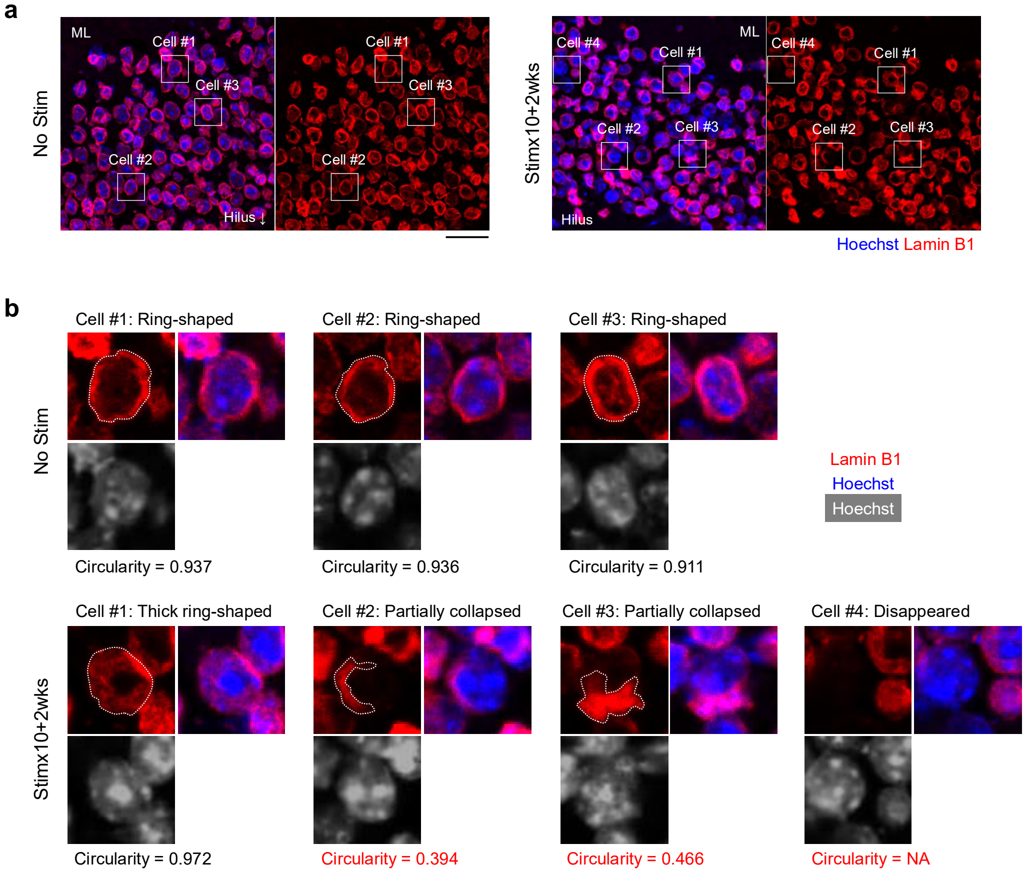


***Supplementary Figure 5. Changes in lamin B1 expression and circularity after REPOPS***

**a,** Lamin B1 (red) and Hoechst (blue) staining in the DG (same data as Fig. 4e). Scale bar, 20 μm.

**b,** Enlarged images of representative neurons (white boxes in **a**). Red, lamin B1; blue/white, Hoechst.

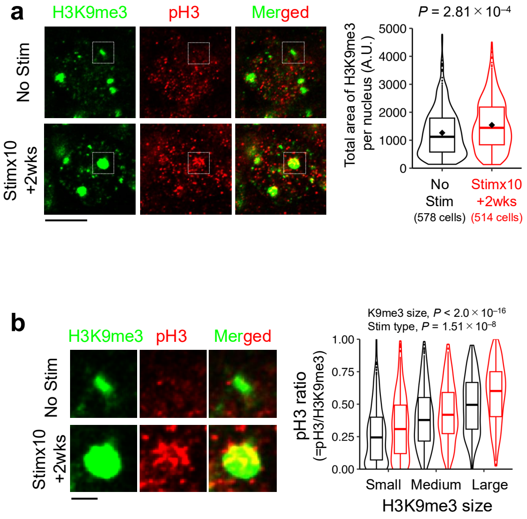


***Supplementary Figure 6. Mitosis-like epigenetic modifications are associated with heterochromatin enlargement***

**a,** STED microscopy images showing phospho-histone H3 (pH3; red) and histone H3 lysine 9 tri-methylation (H3K9me3; green), a marker of highly condensed heterochromatin domains[3]. Scale bar, 5 μm. The violin-box plot shows the size distribution of H3K9me3-positive regions. Centre lines represent medians; box boundaries show upper and lower quartiles; whiskers represent maxima and minima; and diamonds denote mean values. 578 and 514 neurons from four biologically independent samples in the No Stim and Stim×10+2wks groups, respectively. Welch's *t*-test, $t_{(1018)} = 5.32$, $P = 1.25 \times 10^{-7}$.

**b,** Enlarged views of the white boxed regions in **a**. Scale bar, 1 μm. The box plot shows the ratio of pH3 signal coverage per each H3K9me3 dot (i.e., the area of $pH3^{+}H3K9me3^{+}$ double-positive regions normalized by the H3K9me3 dot size). H3K9me3 dots were categorized into Small (< 500 A.U.), Medium (500–1000 A.U.), and Large (>1000 A.U.). Two-way ANOVA: H3K9me3 size, $F_{(2, 2171)} = 186.8$, $P < 2.0 \times 10^{-16}$; stimulation type, $F_{(1, 2171)} = 32.3$, $P = 1.51 \times 10^{-8}$, interaction, $F_{(2, 2171)} = 1.513$, $P = 0.22$.

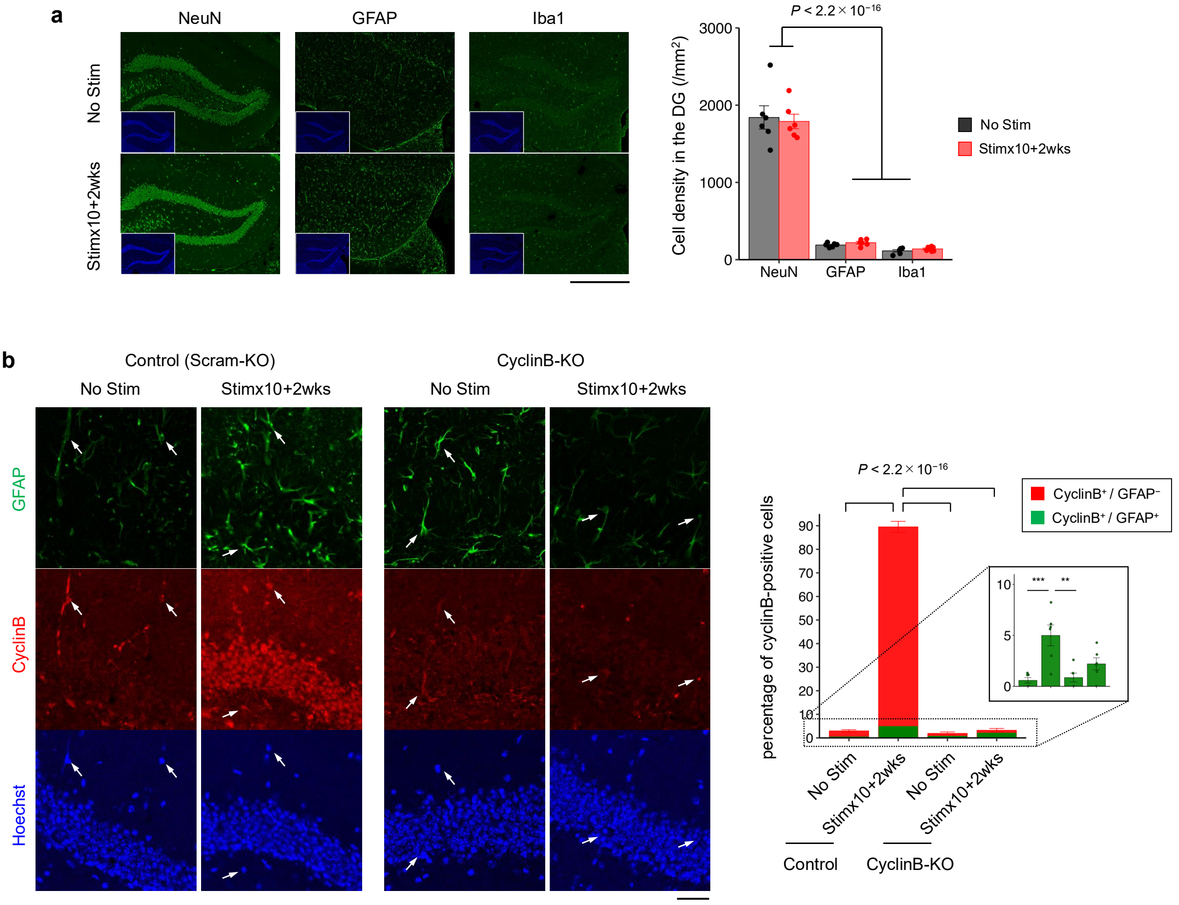


***Supplementary Figure 7. Limited contribution of glial cells to REPOPS-induced cyclin B expression***

**a,** Immunostaining for cell-type markers (NeuN, excitatory neurons; GFAP, astrocytes; Iba1, microglia) in the DG. Scale bar, 500 µm. Bar graph shows density of cells positive for each marker.

**b,** Confocal images of GFAP and cyclin B co-staining. Arrows indicate GFAP$^+$/cyclin B$^+$ astrocytes. Scale bar, 20 µm. Bar graph showing % of cyclin B$^+$ cells among Hoechst$^+$ cells (left), and GFAP$^+$/cyclin B$^+$ cells (right). One-way ANOVA; $F_{(3,20)} = 1092$ and $F_{(3,20)} = 24.4$, $P < 2.0 \times 10^{-16}$ and $P = 3.45 \times 10^{-4}$. Bonferroni correction was applied for multiple comparisons. $^{**}P < 0.01$, $^{***}P < 0.001$.

REPOPS increased the number of cyclin B$^+$ cells in the Scram-KO group. Although GFAP$^+$/cyclin B$^+$ cells were also increased, the majority of cyclin B induction occurred in neurons, indicating limited astrocytic contribution.

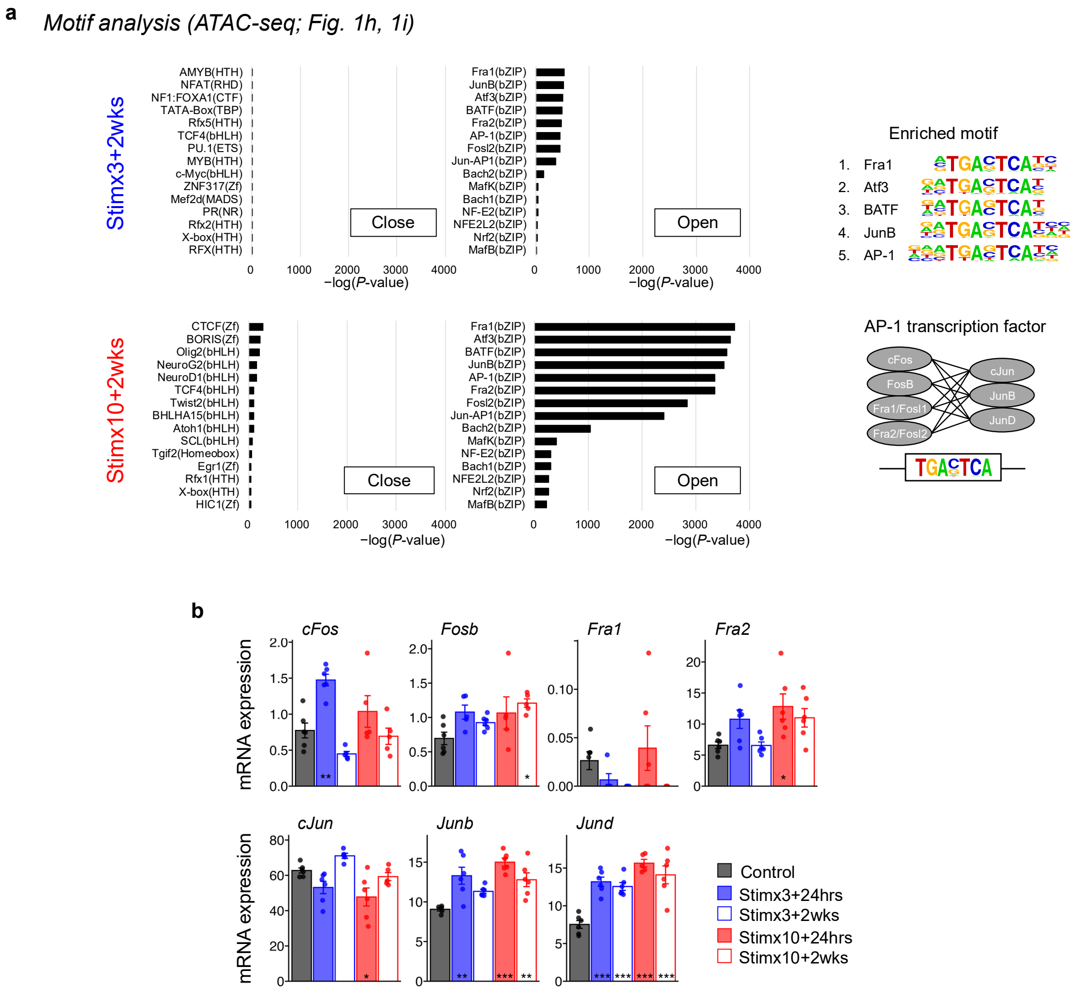


***Supplementary Figure 8. AP-1 motifs are enriched in chromatin regions opened by REPOPS***

**a,** Motif enrichment analysis of ATAC-seq peaks (Fig. 1h, 1i) performed using HOMER. The bar graph shows –$\log_{10}$($P$-value) for similarity between altered chromatin regions in ATAC-seq data and known ChIP-seq profiles. The enriched motifs of the top five in Stim×10+2wks group are shown on the right.

**b,** Expression of AP-1 genes. Data are shown as mean ± s.e.m. (n = 6 mice/group). One-way ANOVA with Bonferroni correction; *$P$ < 0.05, **$P$ < 0.01, ***$P$ < 0.001.

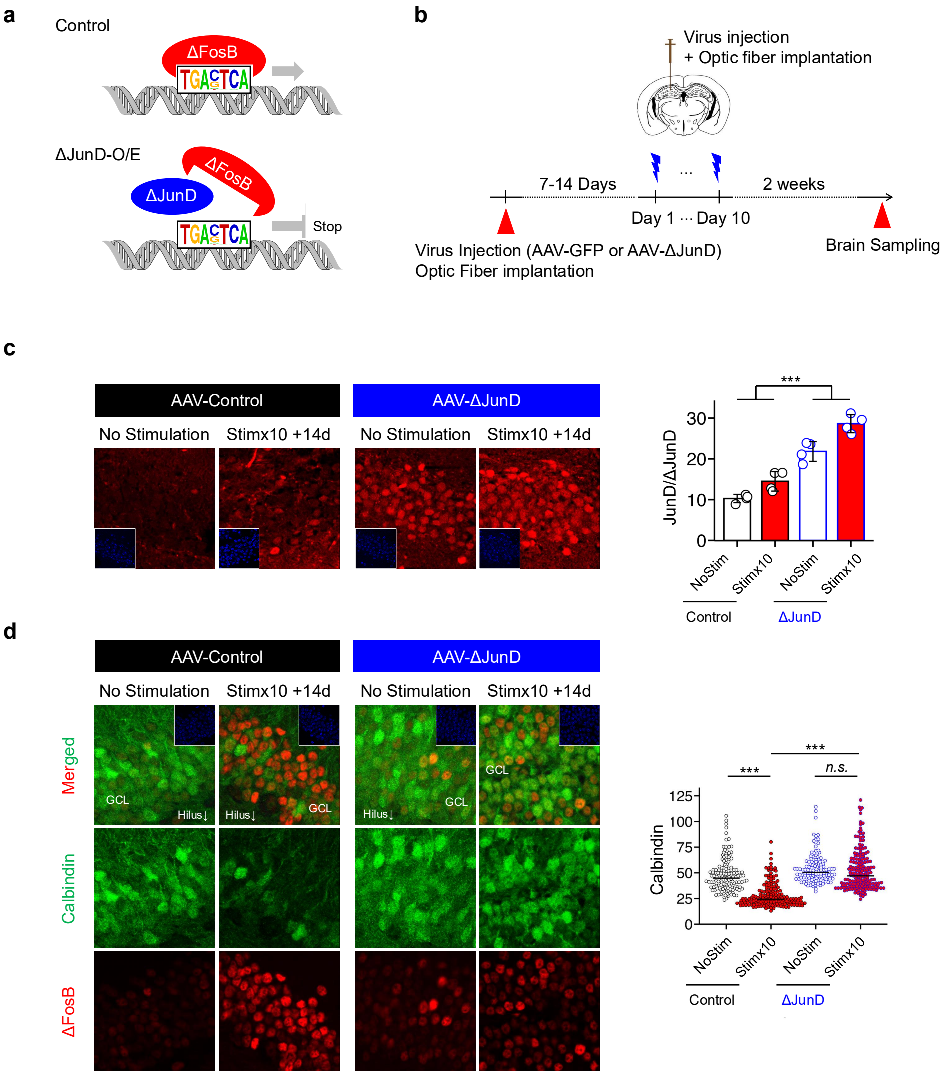


***Supplementary Figure 9. ΔFosB reduces calbindin expression in DG neurons.***

**a,** ΔJunD: mutant form of JunD that retains DNA-binding ability but lacks transactivation potential, acting as a dominant-negative inhibitor of AP-1 by forming non-functional dimers with AP-1 family members.

**b,** Schematic of the experimental design.

**c,** Immunostaining of JunD/ΔJunD in the DG. Bar graph shows JunD/ΔJunD immunoreactivity. n = 4 mice per group. Data are mean ± s.e.m. One-way ANOVA: $F_{(3, 12)} = 59.56$, $P = 1.77 \times 10^{-7}$; post hoc comparisons using Tukey's test, ***$P < 0.001$.

**d,** Immunostaining of calbindin and ΔFosB in the DG. Beeswarm plot shows calbindin immunoreactivity in individual neurons (n = 142-182 neurons per group). Black bars indicate the median. One-way ANOVA: $F_{(3, 645)} = 100.63$, $P < 2.0 \times 10^{-16}$; post hoc comparisons using Tukey's test, ***$P < 0.001$.

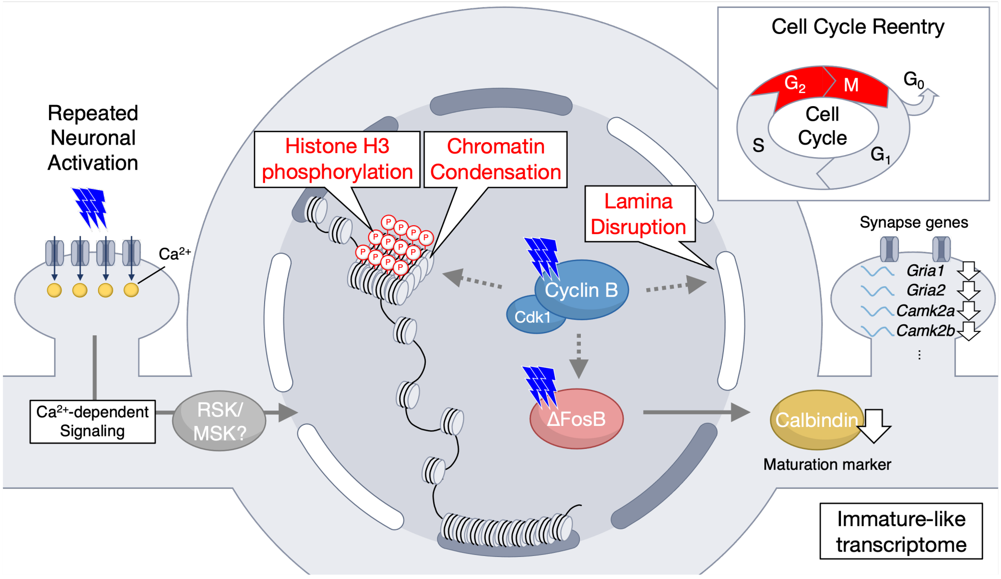


***Supplementary Figure 10. Nuclear reprogramming: molecular mechanism of dematuration induced by repeated neuronal activation.***

Repeated neuronal activation leads to $Ca^{2+}$ influx and subsequent activation of intracellular signalling pathways, inducing a $G_2$/M-like transcriptional program. Activation of Ccnb/Cdk1, a key driver of the $G_2$/M-phase transition, triggers nuclear reprogramming, including nuclear lamina disruption, histone H3 phosphorylation, and chromatin condensation. Repeated activation also induces the expression of AP-1 transcription factors, particularly ΔFosB. Under conditions of sustained Ccnb/Cdk1 activity, ΔFosB expression is stably maintained at elevated levels and leads to an immature-like transcriptional signature, including decreased expression of calbindin, a marker of neuronal maturation. These processes may underlie activity-dependent alterations in neural information coding and contribute to the antidepressant effects of brain stimulation therapies.

***Supplementary Table 1. Statistical Information of behavioural tests***

### *Supplementary Result*

#### *Supplementary result 1. Essential role of $Ca^{2+}$ influx and kinase activation in the induction of a cellular immature state (Related to Supplementary Fig. 2)*

To identify the signaling pathways underlying REPOPS-dependent dematuration, we developed the "Optfluidic neural probe," which enables simultaneous drug delivery and optogenetic stimulation within the same brain region via an implanted cannula (Supplementary Fig. 2a). Using this device, we administered a drug followed by 5 minutes of optogenetic stimulation daily for 10 consecutive days (Supplementary Fig. 2b). Brain tissue was collected 24 hours after the final stimulation, and the effect of the drug on dematuration was evaluated by assessing calbindin expression levels.

To investigate whether $Ca^{2+}$ influx is essential for dematuration, we performed REPOPS with administration of a cocktail of $Ca^{2+}$ blockers, including NBQX (AMPA-R antagonist), AP5 (NMDA-R antagonist), mibefradil (non-specific blocker of voltage-gated $Ca^{2+}$ channels), and nimodipine ($Ca_V1$ blocker). In the saline-treated control group, REPOPS significantly reduced calbindin expression (Supplementary Fig. 2c). In contrast, calbindin expression was unchanged in the $Ca^{2+}$ blocker-treated group (Supplementary Fig. 2c). These results indicate that $Ca^{2+}$ influx is crucial for neuronal dematuration. Notably, these findings also rule out the possibility that dematuration is merely an artifact of REPOPS (e.g., heat-induced tissue damage or inflammation). In line with this, it was further confirmed in Supplementary Fig. 1d that dematuration was not an artifact caused by surgery-induced inflammation or heat shock from optogenetic stimulation.

To explore the kinases involved in dematuration, we conducted REPOPS with administration of four kinase inhibitors with distinct inhibition spectra: K252a, SB218078, Dasatinib, and Ro-32-0432. K252a, SB218078, and Ro-32-0432 successfully suppressed dematuration, whereas Dasatinib did not (Supplementary Fig. 2d, 2e). Based on these results, we screened candidate kinases potentially involved in dematuration. Using a comprehensive pairwise kinase-inhibitor activity dataset[1], we compared the inhibition spectra of the effective inhibitors with that of the ineffective inhibitor (Supplementary Fig. 2f). Kinases that were more strongly inhibited by the effective inhibitors (K252a, SB218078, and Ro-32-0432) than the ineffective inhibitor (Dasatinib) were shortlisted as candidates. This analysis identified 57 potential kinases, of which the top 25 are shown in Supplementary Fig. 2f. This candidate group includes RSK/MSK, which are known to regulate chromatin remodeling via histone phosphorylation[4,5], and CDK, a key regulator of the cell cycle. These findings suggest that the activation of $Ca^{2+}$-dependent phosphorylation signaling pathways plays a pivotal role in

REPOPS-dependent dematuration.

***Supplementary result 2. Behavioral analysis of optogenetically-stimulated mice (Related to Supplementary Fig. 3)***

We assessed the impact of REPOPS on memory function, which can be impaired in ECT patients by conducting contextual and cued fear-conditioning tests (Supplementary Fig. 3a). In the conditioning session, there was a significant effect of stimulation type on the distance travelled ($P = 5.43 \times 10^{-5}$), but freezing time did not differ significantly difference in ($P = 0.631$; Supplementary Fig. 3a (i)). In the context test 24 hours after the conditioning test, there was no significant effect of stimulation type on the distance travelled or freezing, but mice in the Stim×10 group exhibited a shorter freezing time than No Stim groups in the first minutes ($P = 0.0394$; Supplementary Fig. 3a (ii)). In the cued test with different contexts, there was no significant effect of stimulation type on the distance travelled and freezing (Supplementary Fig. 3a (iii)). In the context test one month after the conditioning, there was a significant effect of stimulation type on the distance travelled ($P = 0.006$), and mice in the Stim×10 group traveled significantly longer distance than No Stim groups in the 1–4 minutes ($P < 0.05$; Supplementary Fig. 3a (iv)). Mice in the Stim×10 group exhibited a significantly shorter freezing time than No Stim groups in the last minutes ($P = 0.0223$). In the cued test one month after the conditioning test, there was no significant effect of the stimulation type on the distance travelled and freezing time (Supplementary Fig. 3a (v)). Together, these data suggest that REPOPS impairs contextual memory, consistent with known ECT side effects.

We assessed the impact of REPOPS on sociability using a social interaction test (Supplementary Fig. 3b). Mice in the No Stim and Stim×3 groups spent significantly more time in the chamber with stranger mice than in the chamber with empty cages (Supplementary Fig. 3b (i)). In contrast, no significant difference was observed in the time spent in the stranger and empty chambers in the Stim×10 group (Supplementary Fig. 3b (i)). A similar trend was observed for time spent around the cages (Supplementary Fig. 3b (ii)). The preference index (i.e., the ratio of time spent in the stranger cage to the total time) was significantly smaller in Stim×10 group than that in No Stim group (P = 0.034; Supplementary Fig. 3b (iii)). These observations suggest that REPOPS reduces a preference for social novelty in mice.
